# The head direction circuit of two insect species

**DOI:** 10.1101/854521

**Authors:** Ioannis Pisokas, Stanley Heinze, Barbara Webb

## Abstract

Recent studies of the Central Complex in the brain of the fruit fly have identified neurons with activity that tracks the animal’s heading direction. These neurons are part of a neuronal circuit with dynamics resembling those of a ring attractor. Other insects have a homologous circuit sharing a generally similar topographic structure but with significant structural and connectivity differences. We model the connectivity patterns in two insect species to investigate the effect of the differences on the dynamics of the circuit. We illustrate that the circuit found in locusts can also operate as a ring attractor and identify differences that enable the fruit fly circuit to respond faster to heading changes while they render the locust circuit more tolerant to noise. Our findings demonstrate that subtle differences in neuronal projection patterns can have a significant effect on the circuit performance and emphasise the need for a comparative approach in neuroscience.

## Introduction

For a variety of behaviours that relocate an insect in its environment, it is important for the animal to be able to keep track of its heading relative to salient external objects. This external reference object could be a nearby target, a distant landmark or even a celestial beacon. In insects, the discovery of a neuronal circuit with activity that tracks heading direction provides a potential basis for an internal compass mechanism (***Zhang, 1996***; ***Homberg, 2004***; ***Heinze and Reppert, 2012***). Such an internal compass can mediate a simple navigation competence such as maintaining a straight course (***Dacke et al., 2003; Mouritsen and Frost, 2002***) or reorienting to a target after distractions (***Neuser et al., 2008***), but is also essential for the more complex navigational process of path integration (or dead reckoning) which enables central-place foragers to return directly to their nest after long and convoluted outward paths (***Darwin, 1873***; ***von Frisch, 1967***; ***Mittelstaedt and Mittelstaedt, 1980***; ***Müller and Wehner, 1988***). While the neural basis underlying these navigation strategies are not known in detail, a brain region called the central complex (CX) is implicated in many navigation related processes.

The CX of the insect brain is an unpaired, midline-spanning set of neuropils that consist of the protocerebral bridge (PB), the ellipsoid body (also called lower division of the central body), the fan-shaped body (also called upper division of the central body) and the paired noduli. These neuropils and their characteristic internal organisation in vertical slices, combined with horizontal layers are highly conserved across insects. This regular neuroarchitecture is generated by sets of columnar cells, innervating individual slices, as well as large tangential neurons, innervating entire layers. The structured projection patterns of columnar cells result in the PB being organised in 16 or 18 contiguous glomeruli and the ellipsoid body (EB) in 8 adjoined tiles.

Crucially, the CX is of key importance for the computations required to derive a heading signal (***Pfeiffer and Homberg, 2013***; ***Triphan et al., 2010***; ***Neuser et al., 2008***; ***Ofstad et al., 2011***; ***Homberg, 2004***; ***Homberg et al., 2011***). In locusts (*Schistocerca gregaria*), intracellular recordings have revealed a neuronal layout that topographically maps the animal’s orientation relative to simulated skylight cues, including polarized light and point sources of light (***Heinze and Homberg, 2007***; ***el Jundi et al., 2014***; ***Pegel et al., 2019***). Calcium imaging of columnar neurons connecting the EB and the PB (E-PG neurons) in the fruit fly *Drosophila melanogaster*, revealed that the E-PG neuronal ensemble maintains localised spiking activity — commonly called an activity ‘bump’ — that moves from one group of neurons to the next as the animal rotates with respect to its surrounding (***Seelig and Jayaraman, 2015***; ***Giraldo et al., 2018***). This has been confirmed for restrained flies walking on an air-supported rotating ball (***Seelig and Jayaraman, 2015***) as well as tethered flies flying in a virtual reality environment (***Kim et al., 2017***). Notably, the heading signal (the activity ‘bump’) is maintained even when the visual stimulus is removed, and it moves relative to the (no longer visible) cue as the animal walks in darkness (***Seelig and Jayaraman, 2015***). The underlying circuit therefore combines idiothetic and allothetic information into a coherent heading signal. Overall, this neuronal activity appears to constitute an internal encoding of heading in the insect’s CX, which closely resembles the hypothetical ring attractor (***Amari, 1977***) proposed by ***Skaggs et al.*** (***1995***) to account for the rat ‘head direction’ cells (***Taube et al., 1990***; ***Blair and Sharp, 1995***; ***Redish et al., 1996***; ***Stackman and Taube, 1998***; ***Goodridge et al., 1998***; ***Goodridge and Touretzky, 2000***; ***Sharp et al., 2001***; ***Taube and Bassett, 2003***; ***Stratton et al., 2010***).

In recent years, several computational models of the fly’s CX heading tracking circuit have been presented. Some of these models are abstract while others attempt to ascribe particular roles to neurons (***Cope et al., 2017***; ***Kakaria and de Bivort, 2017***; ***Su et al., 2017***; ***Kim et al., 2017***). ***Cope et al***. (***2017***) proposed a ring attractor model that is inspired by the rat ‘head direction’ cell model of ***Skaggs et al.*** (***1995***). ***Kakaria and de Bivort*** (***2017***) presented a spiking neuronal model consisting of the four types of CX neurons shown to play a role in heading encoding: E-PG, P-EN, P-EG, and Delta7 neurons. Their model demonstrated that these neurons constitute a sufficient set to exhibit ring attractor behaviour. In contrast, ***Su et al.*** (***2017***) implemented a spiking neuronal model consisting of the E-PG, P-EN, and P-EG neurons with inhibition provided by a group of R ring neurons. In both neurobiological studies and computational models the key neurons variously involved in the hypothetical ring attractor circuit are the E-PG, P-EN, P-EG, Delta7 and R ring neurons (***Wolff and Rubin, 2018***; ***Wolff et al., 2015***; ***Kakaria and de Bivort, 2017***; ***Su et al., 2017***; ***Green et al., 2017***; ***Kim et al., 2017***). The E-PG, P-EN and P-EG neurons have been postulated to be excitatory while Delta7 or R ring neurons are conjectured to be mediating the inhibition (***Kakaria and de Bivort, 2017***; ***Su et al., 2017***). The E-PG and P-EN neurons are postulated to form synapses in the PB and in the EB forming a recurrent circuit. The ring attractor state is set by a mapping of the azimuthal position of visual cues to E-PG neurons around the ring which are assumed to be the positional input to this circuit. Furthermore, P-EN neurons shift the heading signal around the ring attractor when stimulated, in a fashion similar to the left-right rotation neurons proposed by ***Skaggs et al.*** (***1995***) (***Turner-Evans et al., 2017***; ***Green et al., 2017***). In principle, two main types of ring attractor implementation exist: One with local excitation and global, uniform, inhibition and another one characterized by sinusoidally modulated inhibition across the ring attractor. ***Kim et al.*** (***2017***) have experimentally explored the type of ring attractor that could underlie the head direction circuit of the fruit fly and concluded that the observed dynamics of E-PG neurons can best be modeled using a ring attractor with local excitation and uniform global inhibition.

The above outlined overall circuit depends critically on the detailed anatomical connections between cell types of the CX, so that the implementation of a specific type of ring attractor imposes additional constraints on the neuronal connection patterns and individual morphologies. Although the CX is highly conserved on a broad level, details at the level of single neurons vary between insect species. Yet, conclusions about the function of the circuit are usually drawn from *Drosophila* data and applied to insects in general. Given numerous differences in the CX neuroarchitecture between insects, we asked whether a ring attractor circuit is also plausible when taking into account anatomical data from another model species, the desert locust.

Three main differences are evident when comparing the CX of the fruit fly and the locust (***Figure 1***). First, as in most insects except *Drosophila*, the EB of the locust is not closed around the edges, but is crescent-shaped, preventing the E-PG neurons from forming a physical ring. Second, the *Drosophila* PB consists of nine glomeruli per hemisphere, and accordingly 18 groups of E-PG neurons. In locusts there are 8 glomeruli per hemisphere and 16 groups of neurons. Third, a key part of the proposed ring attractor circuit, the Delta7 neurons (TB1 neurons in the locust) differ strikingly in their arborization pattern across the width of the PB. Whereas these cells possess two columnar output sites located eight glomeruli apart in all species, their dendrites have an approximately uniform density across the PB glomeruli in *Drosophila*. This differs substantially from the dendritic distribution in the desert locust, in which the postsynaptic domains of the eight Delta7 neurons are restricted to particular glomeruli of the PB, avoiding the regions around the output branches. This pattern is conserved in other species as well, such as in the Monarch butterfly (*Danaus plexippus*), a sweat bee (*Megalopta genalis*), as well as in two species of dung beetles (*Scarabaeus lamarcki* and *Scarabaeus satyrus*) (***Heinze and Homberg, 2007***; ***Heinze et al., 2013***; ***Stone et al., 2017; el Jundi et al., 2018***). As *Drosophila* appears to be different from other insects, we asked which functional consequences can be expected by the homologous neuronal circuits in the different CX varieties and how these functional differences might correlate to behavioural characteristics of each insect.

**Figure 1.**
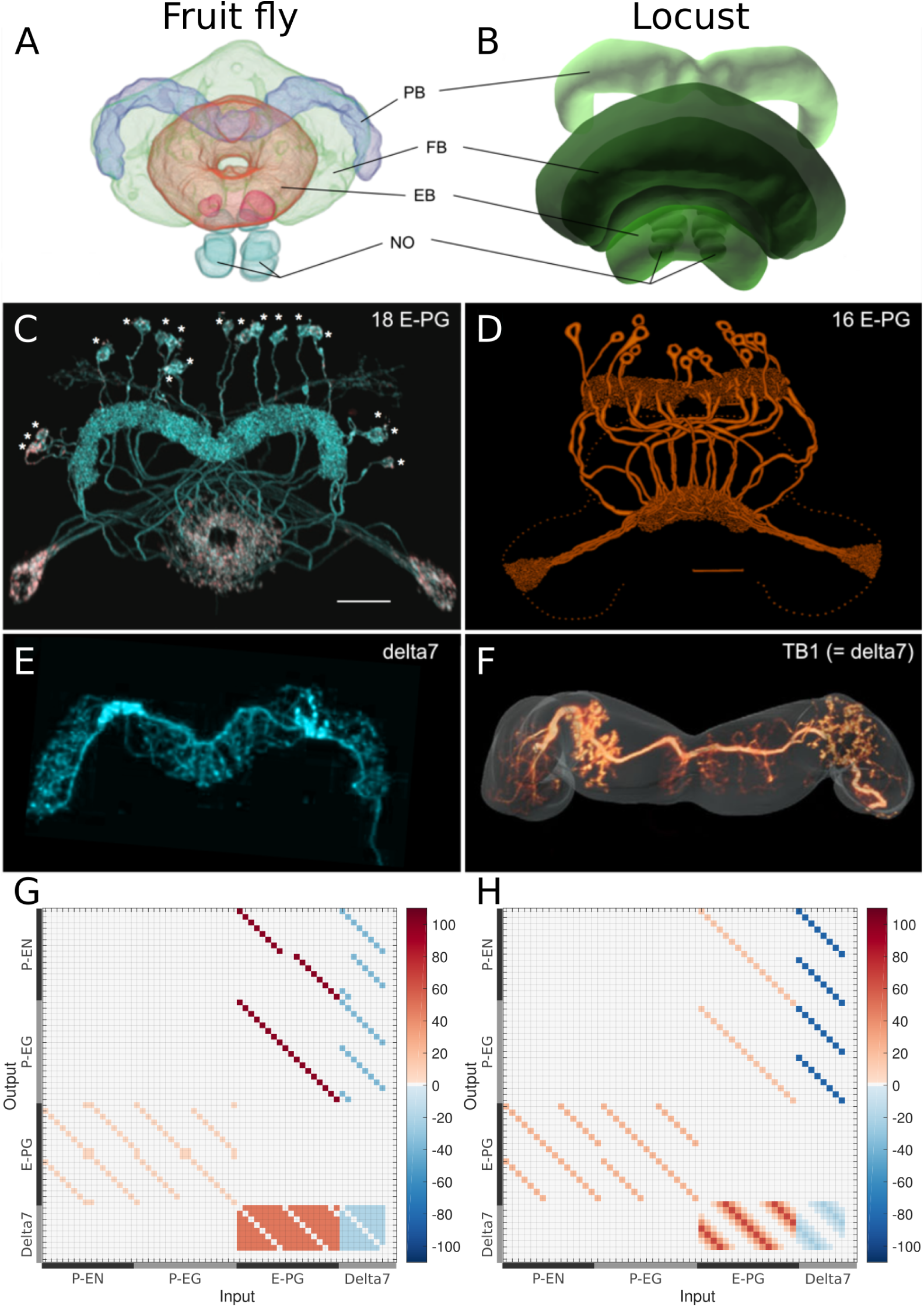
Anatomical differences between two species. There are three apparent differences between the CX of the fruit fly (*Drosophila melanogaster*) and the desert locust (*Schistocerca gregaria*). **A, B**: The ellipsoid body in the fruit fly has a toroidal shape while in the locust is crescent-shaped so its two ends are separate (**A** modified from ***Wolff and Rubin*** (***2018***), **B** Image obtained from Insect Brain Database https://insectbraindb.org, data from ***el Jundi et al.*** (***2010***)). **C, D**: The protocerebral bridge consists of 18 glomeruli and 18 corresponding E-PG (see ***Table 3***) and P-EG neurons in the fruit fly while in the locust there are 16 glomeruli and neurons innervating them (**C** modified from ***Wolff and Rubin*** (***2018***), **D** modified from ***Vitzthum and Homberg*** (***1998***)). **E, F**: The Delta7 neurons in the fruit fly have postsynaptic domains along the whole length of their neurite while in the desert locust only in specific sections with gaps in between (**E** modified from ***Wolff et al.*** (***2015***), **F** modified from ***Beetz et al.*** (***2015***)). **G, H**: The connectivity matrices derived by the exact neuronal projections of the fruit fly (*Drosophila melanogaster*) and the desert locust (*Schistocerca gregaria*), respectively. The difference in the distribution of Delta7 neuron synaptic domains is evident at the lower right part of the images. Synaptic strength derived by an optimisation process, as described in Methods and Materials, is denoted by colour.

To explore this question, we have used the anatomical projection patterns of the main CX neuron types in flies and locusts and derived the effective neuronal circuits by simplifying anatomical redundancy. Both resulting circuits indeed have the structural topology of a ring attractor. Despite significant anatomical differences the homologous circuits in the fruit fly and the locust are structurally very similar but not identical. Their differences have significant functional effect in the ability of the two circuits to track fast rotational movements and to maintain a stable heading signal. Our results highlight that even seemingly small differences in the distribution of dendritic fibers can affect the behavioural repertoire of an animal. These differences, emerging from morphologically distinct single neurons, raise the question of how valid broad generalisations from data derived of individual species such as *Drosophila melanogaster* are and highlight the importance of a comparative approach to neuroscience.

**Table 1.** Characteristics of the neuron tuning curves. The Full Width at Half Maximum (FWHM), the peak impulse rate of each family of neurons and the activity amplitude measured as the range of firing rates are shown. Numbers are given as median and standard deviation. The activity of Delta7 neurons in *Drosophila* is approximately even, hence the corresponding FWHM measurement is not meaningful and marked as ‘N/A’.

| Neuron Class | *Drosophila* |  |  | Locust |  |  |
| --- | --- | --- | --- | --- | --- | --- |
|  | FWHM (°) | Peak (imp./s) | Amplitude (imp./s) | FWHM (°) | Peak (imp./s) | Amplitude (imp./s) |
| E-PG | 94.7 ±4.0 | 208.4 ±2.3 | 208.2 ±2.2 | 73.4 ±2.6 | 220.8 ±1.4 | 220.8 ±1.4 |
| P-EN | 74.6 ±3.8 | 230.3 ±2.3 | 230.3 ±2.3 | 58.9 ±3.1 | 163.6 ±0.9 | 163.6 ±0.9 |
| P-EG | 74.6 ±3.8 | 230.3 ±2.3 | 230.3 ±2.3 | 58.9 ±3.1 | 163.6 ±0.9 | 163.6 ±0.9 |
| Delta7 | N/A | 289.9 ±1.8 | 58.1 ±4.2 | 96.0 ±3.2 | 265.4 ±2.9 | 265.4 ±2.9 |

**Table 2.** Characteristics of the activity ‘bump’. The Full Width at Half Maximum (FWHM), the peak impulse rate of the activity ‘bump’ formed across each family of neurons and the amplitude of the activity ‘bump’ measured as the range of firing rates are shown. Measurements were made 10s after stimulus was removed. Numbers are given as median and standard deviation. The activity of Delta7 neurons in *Drosophila* is approximately even, hence the corresponding FWHM measurement is not meaningful and marked as ‘N/A’.

| Neuron Class | *Drosophila* |  |  | Locust |  |  |
| --- | --- | --- | --- | --- | --- | --- |
|  | FWHM (°) | Peak (imp./s) | Amplitude (imp./s) | FWHM (°) | Peak (imp./s) | Amplitude (imp./s) |
| E-PG | 88.3 ±0.3 | 161.0 ±0.2 | 160.1 ±0.3 | 68.3 ±0.1 | 192.6 ±0.1 | 192.0 ±0.2 |
| P-EN | 80.4 ±0.4 | 190.1 ±0.2 | 190.1 ±0.2 | 63.1 ±0.3 | 153.5 ±0.1 | 153.5 ±0.1 |
| P-EG | 71.0 ±0.2 | 190.1 ±0.2 | 190.1 ±0.2 | 63.1 ±0.3 | 153.5 ±0.1 | 153.5 ±0.1 |
| Delta7 | N/A | 274.7 ±0.1 | 27.1 ±0.2 | 101.1 ±0.2 | 266.6 ±0.2 | 266.6 ±0.2 |

**Table 3.** Neuronal nomenclature. The names used for the homologous neurons differ between *Drosophila* and other species. The first column shows the name used in the current paper to refer to each group of neurons. The other three columns provide the names used in the literature.

| Model | *Drosophila* |  | Locust |
| --- | --- | --- | --- |
| Neuron name | Consensus name | Systematic name (Wolff et al., 2018) | Name |
| E-PG | E-PG and E-PG<sub>T</sub> | PBG1–8.b-EBw.s-D/V GA.b and PBG9.b-EB.P.s-GA-t.b | CL1a |
| P-EN | P-EN | PBG2–9.s-EBt.b-NO1.b | CL2 |
| P-EG | P-EG | PBG1–9.s-EBt.b-D/V GA.b | CL1b |
| Delta7 | Delta7 | PB18.s-GxΔ7Gy.b and PB18.s-9i1i8c.b | TB1 |

## Results

### The effective circuit

The neuronal projection data of the fruit fly and the desert locust were encoded in connectivity matrices and used for the simulations we report here (***Figure 1***). While some simplifications could not be avoided, we have exclusively used the anatomically veri1ed projection patterns for each species to construct the connectivity matrices. Since connectivity matrices are not amenable to facilitating conceptual understanding of the underlying circuit, we here analysed the effective connectivity of these neuronal components of the CX for both species.

#### Inhibitory circuit

First, we focus on the inhibitory portion of the circuit. Study of the actual neuronal anatomy of Delta7 neurons in the PB shows that, in both species, each Delta7 neuron has presynaptic terminal domains in two or three glomeruli along the PB (***Heinze and Homberg, 2007***; ***Wolff and Rubin, 2018***). These presynaptic terminal domains are separated by seven glomeruli (***Figure 2***A and 2D). In *Drosophila* the Delta7 neurons have postsynaptic terminals across all remaining glomeruli of the PB (***Wolff and Rubin, 2018***; ***Franconville et al., 2018***) while in locusts the Delta7 neurons have postsynaptic terminal domains only in specific glomeruli (***Heinze and Homberg, 2007***; ***Beetz et al., 2015***).

**Figure 2.**
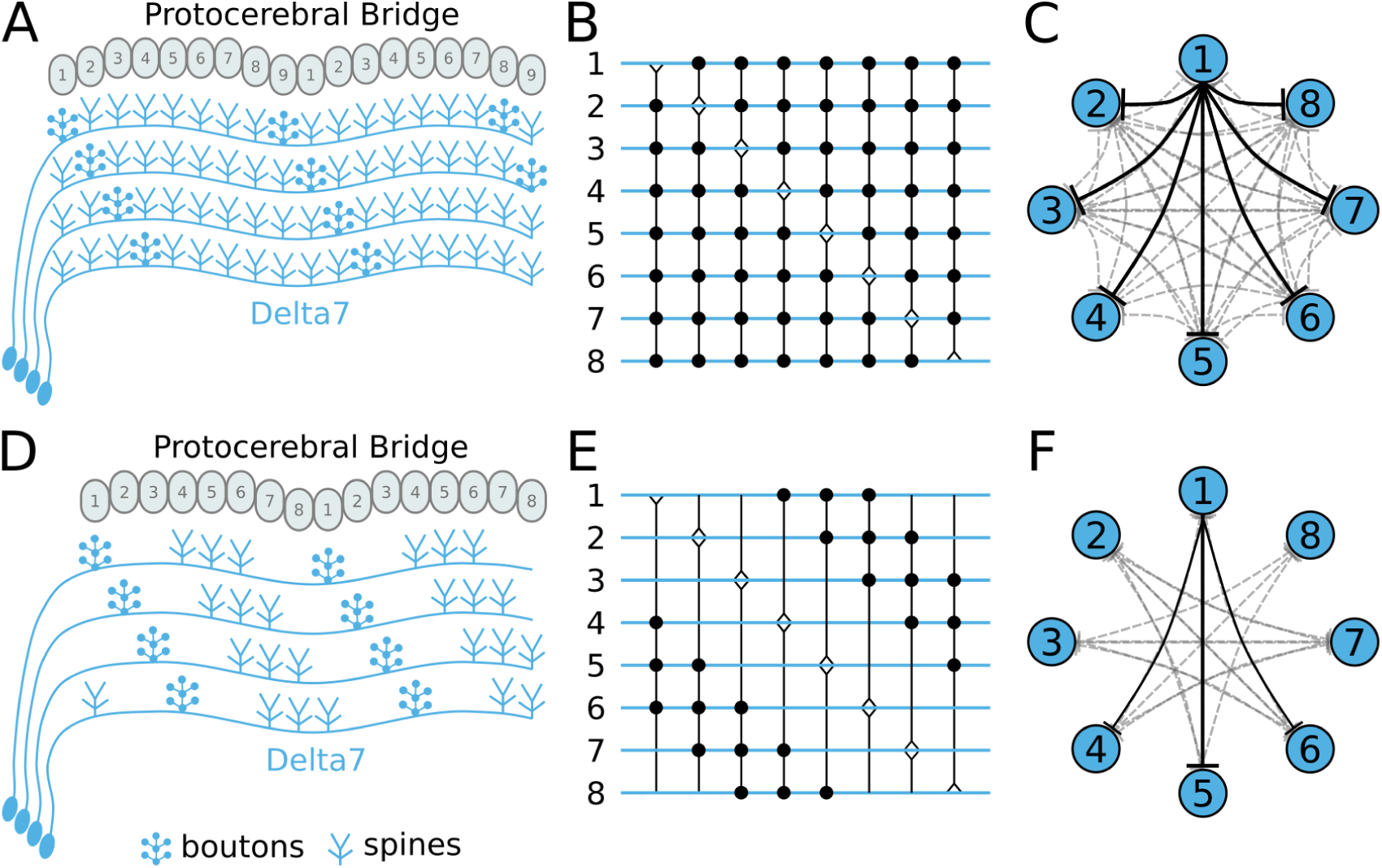
Effective connectivity of the inhibitory (Delta7) neurons. On the top row is the fruit fly circuit, on the bottom row is the locust circuit. In **A** and **D**, four examples of how the eight types of Delta7 neurons innervate the PB are illustrated. In both species, presynaptic domains are separated by seven glomeruli. **B** and **E**: Effective connectivity. Each horizontal blue line represents one Delta7 neuron. Vertical lines represent axons, with triangles indicating outputs from Delta7 neurons and filled circles representing inhibitory synapses between axons and other Delta7 neurons. **C** and **F**: Alternative depiction of the circuit in graph form with blue circles representing Delta7 neurons and lines representing inhibitory synapses between pairs of neurons. Each Delta7 neuron inhibits all other Delta7s in the fruit fly **C**, but only more distant Delta7s in the locust **F**.

There are eight types of Delta7 neurons in the PB, each having the same pattern of synaptic terminals shifted by one glomerulus (Figs 2A and 2D). Within each glomerulus, the Delta7 neuron with presynaptic terminals is assumed to form synapses with all other Delta7 neurons that have postsynaptic terminals in the same glomerulus. Since each Delta7 neuron is presynaptic to the same Delta7 neurons in two or three glomeruli along the PB, we reduce these two or three synaptic domains to one single equivalent synapse between each pair of Delta7 neurons in order to draw a simplified equivalent circuit (Figs 2B and 2E). In order to highlight the main functional differences we redrew these neuronal circuits again in a network graph form which revealed an eight-fold symmetry in both species, regardless the different neuronal anatomies and the anatomical presence of nine PB glomeruli in flies.

The network graph form of the circuit further makes evident a global, uniform, inhibition pattern in the case of the fruit fly versus a local inhibition pattern in the case of the locust (Figs 2C and 2F). That is, in fruit flies each Delta7 neuron forms synapses and inhibits all other Delta7 neurons. On the contrary, in the locust each Delta7 neuron only inhibits a subset of Delta7 neurons with weakening synaptic strengths towards its nearest neighbours. This reveals an effective global inhibition pattern in the fruit fly that fits the finding of ***Kim et al.*** (***2017***) who observed calcium dynamics that better matched a ring attractor with global inhibition in this species.

#### Excitatory circuit

We next focused on the excitatory portion of the hypothetical ring attractor circuit. For deriving the effective circuit of the excitatory portion of the network it was necessary to employ an unconventional numbering scheme for the PB glomeruli; that is, in both hemispheres glomeruli are numbered incrementally from left to right, 1-9 for the fruit fly (***Figure 3***) and 1-8 for the locust (***Figure 4***). EB tiles were numbered 1 to 8 for both species. For brevity, throughout this text we denote a tile numbered 1 as T1 and a glomerulus numbered 1 as G1. Neurons are numbered by the glomerulus they innervate, using a numerical subscript, e.g. P-EN_1_ for the P-EN neurons innervating glomeruli G1.

**Figure 3.**
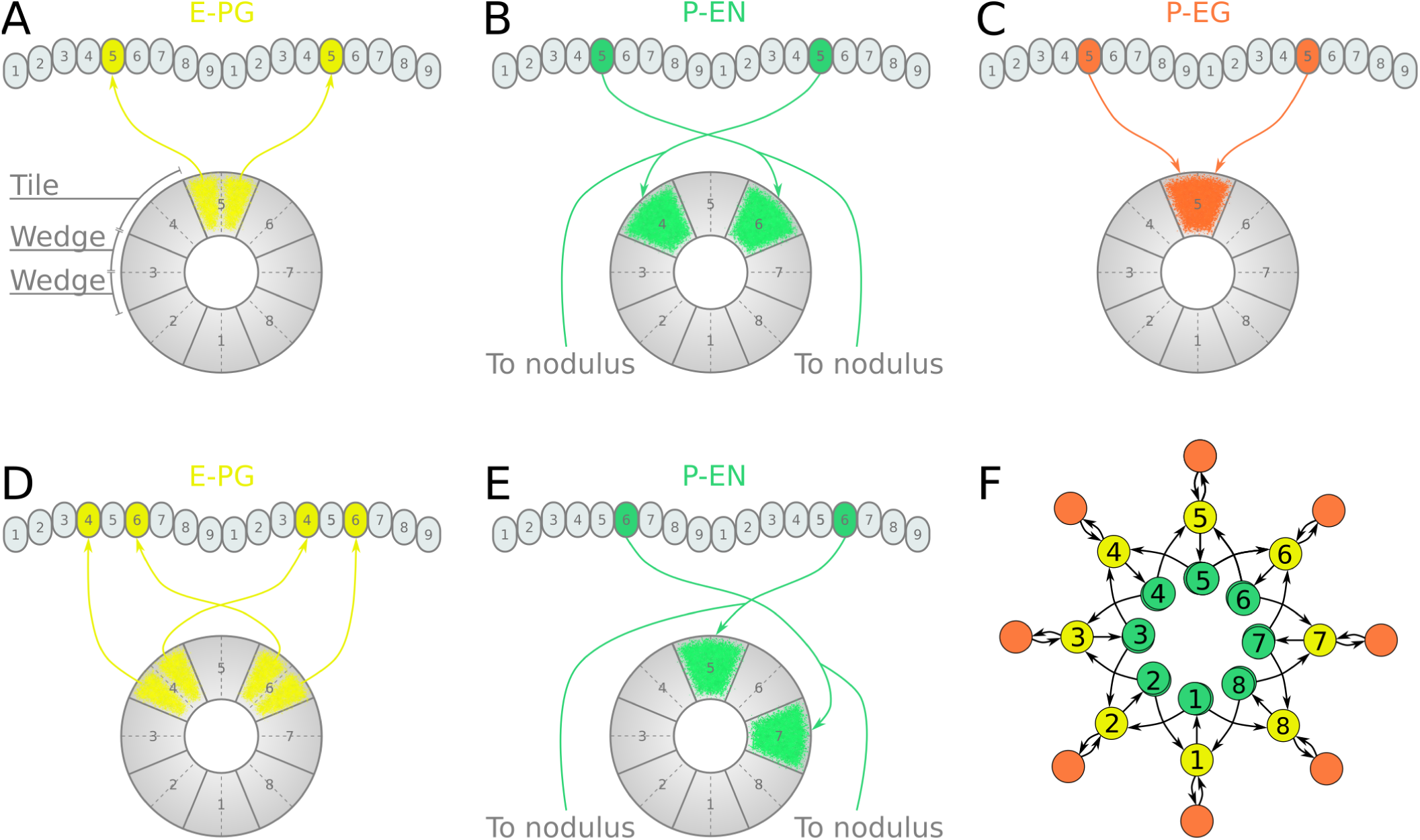
Neuronal projections and effective connectivity in the fruit fly. Projection patterns of the excitatory portion of the fruit fly circuit. **A**–**E**: Examples of E-PG (combined E-PG and E-PG_*T*_, see ***Table 3***), P-EN and P-EG neurons with their synaptic domains and projection patterns (see main text for detailed description). **F**: Conceptual depiction of the effective connectivity of the ring attractor circuit. Each coloured circle represents a group of neurons with arrows representing excitatory synaptic connections. P-EN neurons are shown overlapped because each receives input only from its contralateral nodulus.

**Figure 4.**
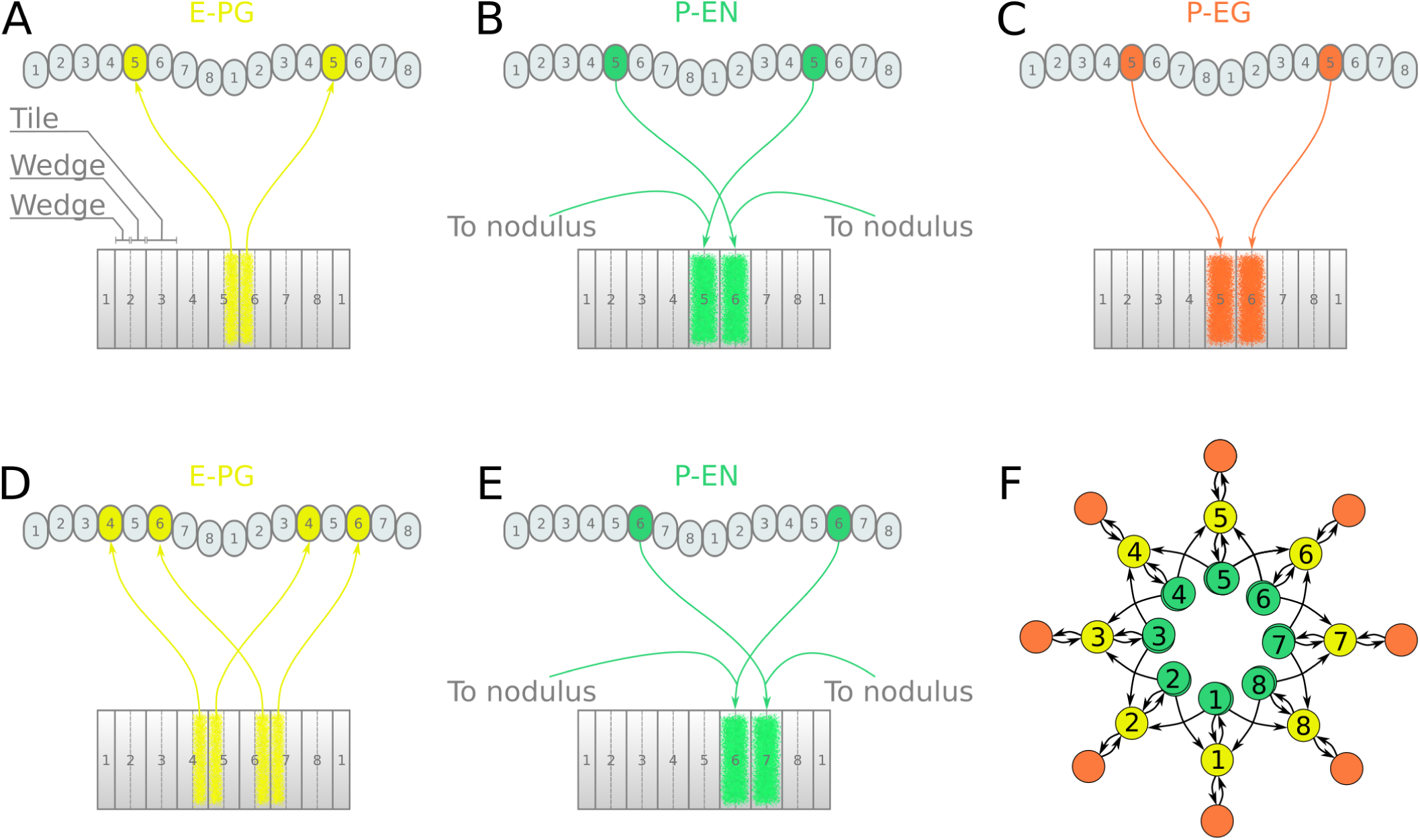
Neuronal projections and effective connectivity in the locust. Projection patterns of the excitatory portion of the locust circuit. **A**–**E**: Examples of E-PG, P-EN and P-EG neurons with their synaptic domains and projection patterns (see main text for detailed description). **F**: Conceptual depiction of the effective connectivity of the ring attractor circuit. Each coloured circle represents a group of neurons with arrows representing excitatory synaptic connections. P-EN neurons are shown overlapped because each receives input only from its contralateral nodulus.

In accordance with calcium imaging (***Turner-Evans et al., 2017***; ***Green et al., 2017***), simulating the fruit fly and locust circuits confirmed that there are two activity ‘bumps’ along the PB. The choice of unconventional numbering scheme for the PB glomeruli has as an effect that both activity ‘bumps’ are centred around neurons innervating identically numbered glomeruli (Supplementary Fig. S1). We used this symmetry to simplify the circuit and derive the effective circuit connectivity.

First, we analyse and derive the effective circuit of the fruit fly. Under our numbering scheme, each E-PG neuron has synaptic domains in identically numbered EB tiles and PB glomeruli (e.g. ***Figure 3***A). That is, E-PG_5_ has synaptic domains in tile T5 and glomeruli G5 in both hemispheres of the PB. P-EN neurons, however, connect corresponding glomeruli from each PB hemisphere to two tiles, one shifted to the left and one to the right, e.g., they would connect glomeruli G5 to tiles T4 and T6 (***Figure 3***B). If we now follow these synaptic pathways one step further, the P-EN_5_ neurons form synapses with E-PG_4_ neurons in T4 and E-PG_6_ neurons in T6, which innervate glomeruli G4 and G6, respectively (***Figure 3***D). These neurons in turn form synapses with P-EN neurons in these glomeruli, making connections back to T5 and onward to T7 (***Figure 3***E). This connectivity pattern continues all the way around the PB glomeruli and EB tiles. Another class of neurons, the P-EG neurons are innervating equally numbered glomeruli and tiles. Hence, they follow the same pattern as the E-PG neurons but with their presynaptic and postsynaptic terminals on opposite ends (***Figure 3***C). Crucially, tile T1 is innervated by both E-PG_1_ and E-PG_9_ which also innervate glomeruli G1 and G9, respectively (Supplementary Fig. S2). This results in tile T1 being innervated by double the number of E-PG neurons than other tiles. But since there are no P-EN neurons innervating the innermost glomeruli (G9 and G1), tiles T2 and T8 are innervated by as many P-EN neurons as any other tile.

By redrawing the circuit in directed graph form we see that surprisingly, the effective circuit of the fruit fly has an eight-fold radial symmetry despite the nine PB glomeruli (illustrated in ***Figure 3***F). To derive this circuit we used the observation that pairs of E-PG_*n*_ neurons connect EB tiles T*n* to PB glomeruli G*n* and since activity is symmetrical in both hemispheres we simplify the circuit by replacing each pair of neurons by one single connection from tile T*n* to glomerulus G*n* (***Figure 3***F). Similarly, due to the symmetrical activation of P-EG_*n*_ neurons innervating equally numbered glomeruli, those pairs of P-EG_*n*_ neurons are also reduced to one unit in the effective circuit (***Figure 3***F). Finally, we preserve two P-EN_*n*_ neurons in each octant, indicated by two overlapped discs in the drawings because even though all P-EN_*n*_ neurons receive the same input in the glomeruli, they also receive differential angular velocity input (***Stone et al., 2017***; ***Turner-Evans et al., 2017***). This equivalent circuit removes the details about the anatomical organisation of the EB and the PB while preserving the effective connectivity.

In the locust the mapping of EB tiles to PB glomeruli is similar to that of the fruit fly (***Figure 4***). Here, E-PG neurons from the two corresponding PB glomeruli in each hemisphere have synaptic domains in two neighbouring EB wedges (half tiles), e.g. E-PG_5_ innervates tiles T5 and T6 an glomeruli G5 of the PB (***Figure 4***A and Supplementary Fig. S3). While pairs of E-PG neurons that innervate identically numbered PB glomeruli receive input from one single tile in the fly, in locusts they receive input from two wedges belonging to two neighbouring tiles. P-EN neurons connect PB glomeruli to tiles shifted by one wedge to the left and right, e.g. glomeruli G5 with tiles T5 and T6 (***Figure 4***B). This is a shift of half tile while in the fruit fly we see a whole tile shift. Finally, P-EG neurons following the same pattern as E-PG neurons, innervate equally numbered glomeruli but two neighbouring tiles, e.g. P-EG_5_ connects G5 to T5 and T6. This is another difference from the fruit fly projection pattern. Tracing this connectivity pattern forward as before (***Figure 4***D and ***Figure 4***E) and redrawing the neurons as circles in a network graph format we get the equivalent network shown in ***Figure 4***F. In spite of the EB in the locust not forming a ring but rather having a crescent shape the effective circuit still forms a ring with an eight-fold symmetry that has structure almost identical to that of the fruit fly.

#### Overall circuit

The similarity between the effective circuits of the locust and the fruit fly was striking. Despite the fact that locusts have eight PB glomeruli while fruit flies have nine, both circuits possessed the same eight-fold symmetry, and the functional role of each neuron class appeared identical. The E-PG neurons were presynaptic to both P-EG and P-EN neurons, with P-EG neurons forming recurrent synapses back to E-PG neurons. P-EN neurons were presynaptic to E-PG neurons with a shift of one octant to the left or right. Overall, two of the main anatomical differences between the two species (eight versus nine PB glomeruli and ring-shaped versus crescent-shaped EB) had no fundamental effect on the principal structure and computations carried out by the CX heading direction circuit. The excitatory portions of the circuits differed only in that the locust P-EN neurons make synapses back to E-PG neurons in the same octant while in the fruit fly they do not. This difference resulted from the P-EN synaptic domains being shifted by half-tile in the locust instead of the whole tile shift seen in the fruit fly. Consequently, the P-EN terminals innervating the middle portion of the two neighbouring synaptic domains in the EB feed back to the same E-PG neurons.

During our analysis of the anatomical data in locusts and flies we also observed that the order of innervation of E-PG neuronal projections in the EB differs between the two species (***Heinze and Homberg, 2008***; ***Williams, 1975***; ***Wolff et al., 2015***; ***Wolff and Rubin, 2018***). Spanning the EB clockwise starting from tile 1, the fruit fly wedges connect first to the right PB hemisphere, then to the left and so on, while in the locust they connect first to the left, then to the right and so on. However, despite this seemingly major difference in projection patterns the effective circuit is preserved between the two species.

When we combined the inhibitory and the excitatory sub-circuits into a complete CX model (Figs 5A and 5B), the E-PG neurons became presynaptically connected to the Delta7 neurons, in line with ***Franconville et al.*** (***2018***). Additionally, each Delta7 neuron inhibits the P-EN and P-EG neurons in the same octant, as well as all other Delta7 neurons (for the fruit fly) or a subset (for the locust), as described above. This difference results to two different types of ring attractor topology; one with global inhibition in the fruit fly and another with local inhibition in the locust.

**Figure 5.**
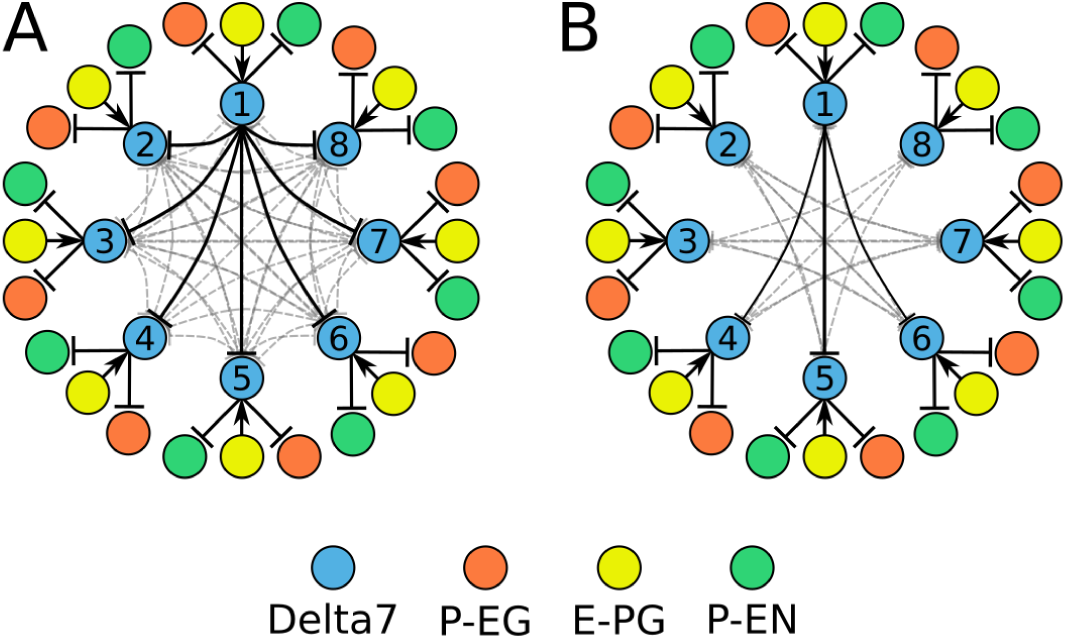
Combined excitatory and inhibitory portion of the ring attractors. Explanatory drawings of the connectivity of the inhibitory portion with the excitatory portion of the circuit for the fruit fly (**A**) and the locust (**B**). Each coloured circle represents one or more neurons with lines representing synaptic connections.

### Predicted synaptic strengths

Assuming that the connectivity we have implemented in our model comprised the necessary and sufficient circuit for a ring attractor in the insect brain, we next investigated what synaptic connectivity strengths were required to produce ring attractor dynamics. This constitutes a prediction for the synaptic strengths we expect to be observed in insects when such measurements become available. To this aim, we ran an optimisation algorithm (described in Methods and Materials) to find regularities in the synaptic pattern sets that resulted in functional ring attractors. A k-means algorithm was used to identify the clusters around which solutions were found. These clusters were ordered by the number of instances found by repetitive runs of the optimiser. Although the absolute synaptic strengths are arbitrary, as they depend on unknown biophysical properties of the involved neurons, a pattern emerged in the relative synaptic strengths between the different synapses (***Figure 6***). The most frequent synaptic strengths patterns were comparably consistent for the fruit fly and the locust. In both species, among the excitatory synapses, the P-EN to E-PG and P-EG to E-PG synaptic strengths were the weakest, while the synaptic strengths from E-PG to P-EG and P-EN neurons were the strongest. The inhibitory synaptic strengths from Delta7 to P-EN and P-EG were stronger in the locust than in the fruit fly, which was consistent with the fly neurons receiving input from more Delta7 neurons.

**Figure 6.**
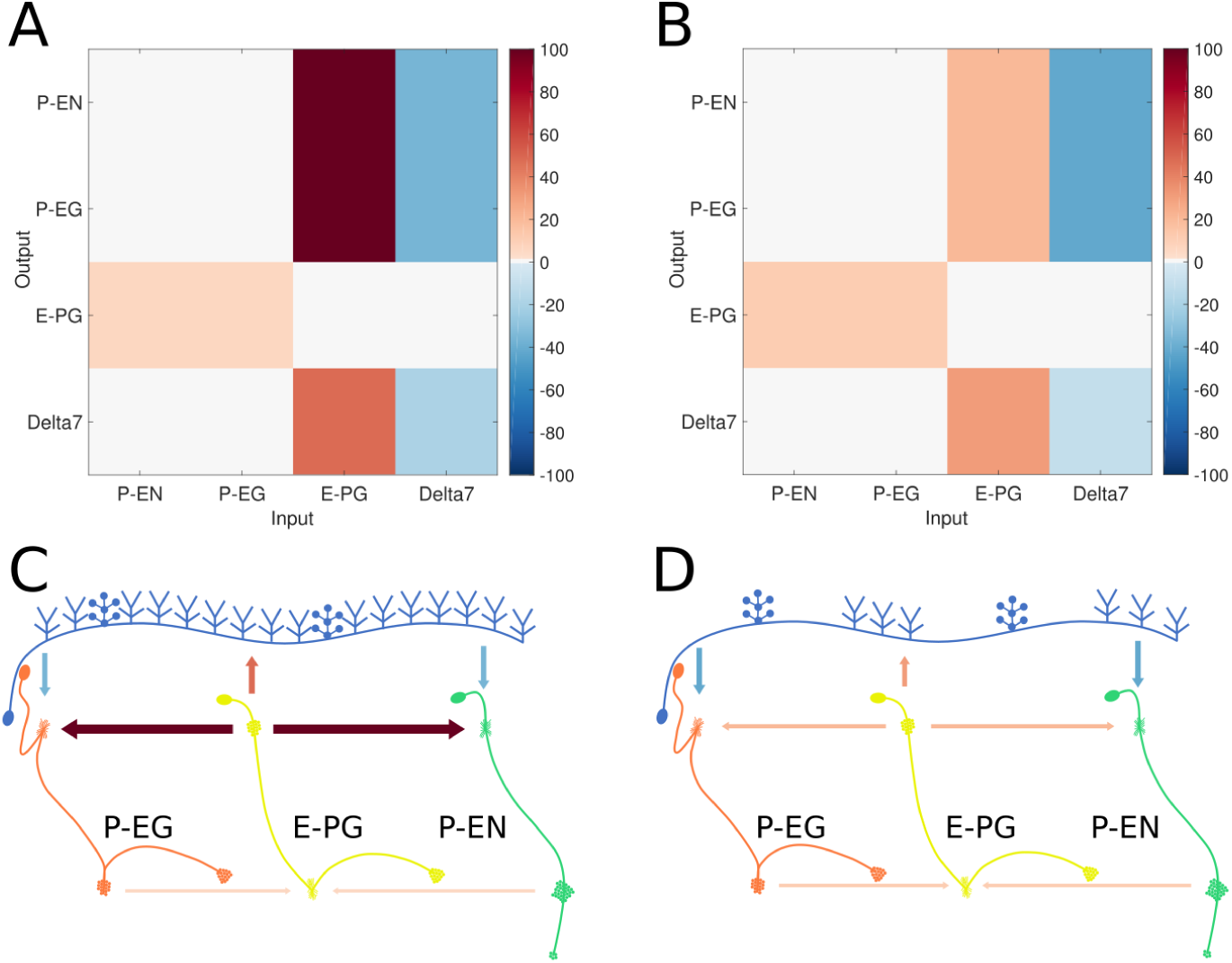
Relative synaptic strengths. Graphical depiction of the synaptic strengths between classes of neurons. **A**,**C**: For the fruit fly ring attractor circuit. **B**,**D**: For the desert locust ring attractor circuit. Synaptic strengths are denoted by colour in panels **A** and **B**. In panels **C** and **D** synaptic strengths between neurons are indicated by arrow colour and thickness in scale. Note that in the locust the synaptic strengths shown for Delta7 neurons are the peak values of the Gaussian distributed strengths shown in ***Figure 1***.

### Predicted neuronal activity

Whereas our simulations confirmed that both the fruit fly and the locust circuit can operate as ring attractors, there were clear differences in the spiking activity and dynamics of the two circuits (***Figure 7***). One major difference was that Delta7 neurons exhibited distinct firing patterns in the two species. In the locust there was a strong heading-dependent modulation in the firing of Delta7 neurons, in line with the heading signal (activity ‘bump’) location. Those Delta7 neurons corresponding to the current heading signal location remained silent. In contrast, in the fruit fly the firing of action potentials was only minimally modulated across the Delta7 population (***Figure 7***A and ***Table 2***). This difference reflected the utilisation of local inhibition in the case of the locust versus the global inhibition in the fruit fly. Electrophysiologists have indeed reported this pronounced firing rate variation in the locust (***Heinze and Homberg, 2007***; ***Heinze et al., 2009***; ***Bockhorst and Homberg, 2015***; ***Pegel et al., 2018***). It will be interesting to see if the fruit fly neurons indeed show a lower modulation as predicted by our model.

**Figure 7.**
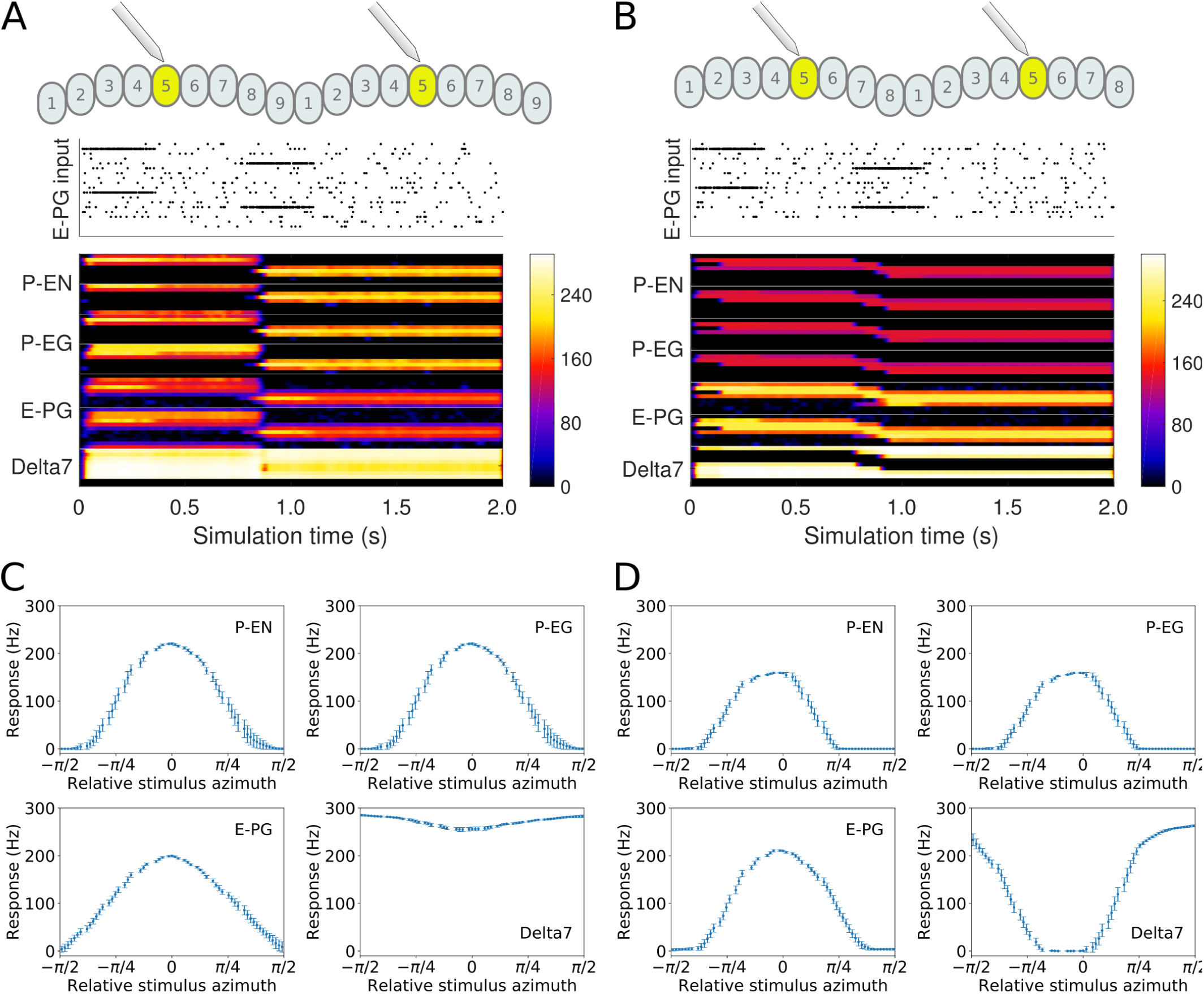
Response to abrupt stimulus changes and tuning curves of neurons. **A** and **B**: The raster plots of the stimuli used to drive the ring attractor during the simulation are shown on top and the spiking rate activity of each neuron at the bottom. In the beginning of the simulation the stimulus spiking activity sets the ring attractor to an initial attractor state. A ‘darkness’ period of no stimulus follows. Then a second stimulus corresponding to a sudden change of heading by 120° is provided. **A**: Response of the fruit fly ring attractor to sudden change of heading. **B**: Response of the locust ring attractor to sudden change of heading. The response of the fruit fly model to sudden changes of heading is faster than that of the locust. **C** and **D**: Response of individual neuron types to different stimuli azimuths (n=40 trials). The mean and standard deviation are indicated by the error bars at the sampled azimuth points. Peak activity has been shifted to 0*°*. **C**: tuning curves for the fruit fly and **D**: tuning curves for the locust.

When comparing the head-direction tuning widths between the two species, we noted that in locusts all cell types are consistently tuned more narrowly (ca. 20%). Within both species, the activity bump is wider for E-PG neurons than for the other excitatory neuron classes (***Table 2***), a difference that is more pronounced in the fruit fly. The tuning of the Delta7 neurons is the widest across cell types in both species (approx. 101° in the locust). In the fruit fly the activity is approximately even across all Delta7 neurons (ca. 10% modulation).

In our models we employed one neuron for each connection, whereas in the actual animals there are multiple copies of each neuron. While definite numbers of neurons will have to await electron microscopical data, there are likely at least two copies of E-PG, P-EG and P-EN neurons in each columnar module, and three to four copies of Delta7 cells (***Williams, 1975***; ***Heinze and Homberg, 2008***; ***Beetz et al., 2015***; ***Wolff et al., 2015***; ***Wolff and Rubin, 2018***). If we were to replace each modeled neuron by a bundle of neurons, the action potential firing rates shown in ***Table 2*** would be divided among the neurons in each bundle. The peak firing rate of each neuron would be in the range of 40–90impulses/s which is similar to the range of the rates recorded electrophysiologically in the locust (***Heinze and Homberg, 2009***).

The steady state peak spiking rate for each group of neurons differs between the fruit fly and the locust circuits. On average, the locust neurons showed ca. 25% higher peak firing rates compared to the fruit fly neurons while the Delta7 neurons have the highest spiking rate in both species. Electrophysiology studies will clarify if this is the case.

The tuning curves of the P-EN and P-EG neurons have the same statistics because in our models we assumed that all neurons have the same biophysical properties and since both these types of neurons receive the same inputs their responses are similar.

### Connectivity differences affect response dynamics

Despite the substantial similarity in functional structure of the two circuits, the subtle differences in connectivity affected the dynamics of the circuit behaviour. This became apparent when we compared the response of both circuits to sudden changes of heading (Figs 7). At a qualitative level, the fruit fly heading signal (the ‘bump’) could jump abruptly from one state to another, whereas the locust circuit exhibited a gradual transition.

To explore whether this difference in movement dynamics of the heading signal could be a result of the different inhibition patterns produced by the Delta7 neurons, we replaced the global Delta7 connectivity pattern in the fruit fly model with the connectivity pattern of the locust Delta7 neurons, effectively swapping the fruit fly version of these cells with the locust version. The data generated by this hybrid-species model revealed that changing the global inhibition to local inhibition was sufficient to produce the gradual ‘bump’ transition we observed in the locust circuit (***Figure 7***B).

#### Quantification of the ring attractor responsiveness

Having shown that small changes in the morphology of the Delta7 cells affect the dynamics of the heading signal in a qualitative way, we next quantified the maximal rate of change each ring attractor circuit could attain. To this end we measured the time it took for the heading signal to transition from one stable location to a new one, in response to different angular heading changes of the stimulus. This was carried out in all three models: the fruit fly model, the locust model, and the hybrid-species model. The fly ring attractor circuit stabilised to the new heading in approximately half the time it takes for the locust circuit to stabilise, across different magnitudes of angular heading change (***Figure 8***A). The hybrid-species circuit had a similar response time to the locust circuit. This confirmed that the pattern of inhibition in the network is the main contributor to the observed effect.

**Figure 8.**
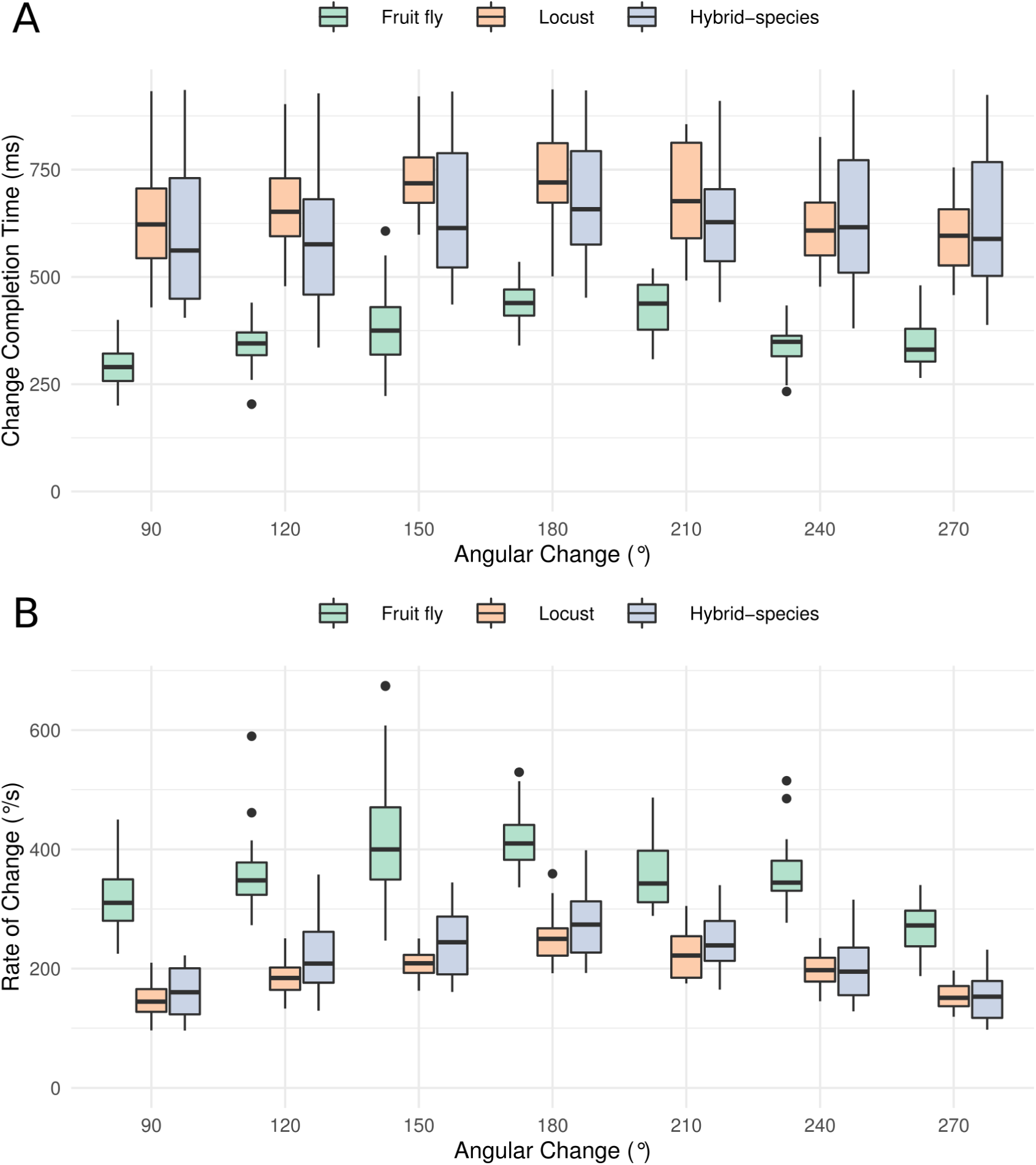
Transition time and rate of the heading signal. **A** Time required from the onset of the stimulus until the heading signal settles to its new state. **B** The maximum rate of angular change each model can attain computed as the ratio of angular change of stimulus divided by transition duration. The values for different magnitudes of heading change are depicted as medians. The boxes indicate the 25^th^ and 75^th^ percentiles. The whiskers indicate the minimum and maximum value in the data after removal of the outliers. ‘Hybrid-species’ is the combination of the fruit fly model with the locust inhibition pattern. Outliers are plotted as black dots.

To calculate the maximal rate of angular change the circuit can possibly track we divided the angular heading change by the time required for the heading signal to transition. Since the heading signal moves along the shortest path around the ring attractor, the numerator is the shortest angular distance calculated as

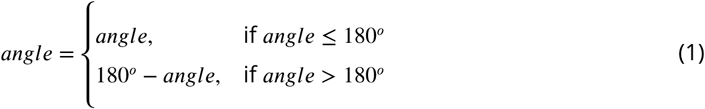

The resulting angular rate of change values revealed that the circuit found in the fruit fly is significantly faster than the locust circuit and the hybrid-species circuit with localised inhibition (***Figure 8***B). The rate of change is maximal for angular displacement of 180°, because the heading signal travels from one azimuth to the next through the shortest path around the ring attractor. Therefore, the maximum azimuth distance it has to possibly travel is 180°. For all other angular displacements there is a shorter than 180° path to the target azimuth. This explains why there is a peak in the angular speed at 180°.

#### Effects of varying the uniformity of inhibition

The above results strongly suggested that the different pattern of inhibition is instrumental to generating the different dynamics in the two circuits. Up to this point we have examined two extreme cases of inhibitory synaptic patterns, that of the global, uniform, inhibition found in *Drosophila* and the localised inhibition found in the locust. However, in principle, there could be any degree of uniformity of the inhibition between these two extremes. So far, the locust inhibition has been modeled as a sinusoid that approximates the synaptic density across the PB glomeruli, estimated by visual inspection of light microscopy data. In the fruit fly, the synaptic distribution of the fruit fly has been modeled as uniform across PB glomeruli, although there might be subtle synaptic density variation along its length. To account for this possibility we explored a range of synaptic domains distributions. As no measurements of synaptic strengths exist for either animal, we asked what effect varying the synaptic terminal distribution has on the ring attractor behaviour. We thus modeled the inhibitory synaptic strength across the PB using two Gaussian functions and varied their variance (*σ*^2^). This would not only give us the effect of different inhibitory widths but would also predict the plausible range of widths that the actual animal must have in order to exhibit the observed dynamics.

Modelling these variations showed that the transition mode of the heading signal depended on both the extent of the inhibitory width and the angular heading change of the stimulus. This sets limits on the plausible variance (*σ*^2^) range that the synaptic strength distribution must obey in the actual animals (***Figure 9***). We observed that for both circuits there was a range of low *σ*^2^ values, corresponding to more localised inhibition, which produce gradual transitions (‘locust-like’). As *σ*^2^ was increased, the inhibitory pattern became more uniform or global, and both circuits transitioned to abrupt jumps (‘fly-like’). The inhibitory synaptic distribution that we inferred visually from light microscopy and used for our initial locust model had value *σ*^2^ = 0.4, corresponding to the gradual activity transition regime across the whole range of angular changes. These results suggested that the pattern of inhibition is indeed key to the circuit dynamics in response to rapid heading changes.

**Figure 9.**
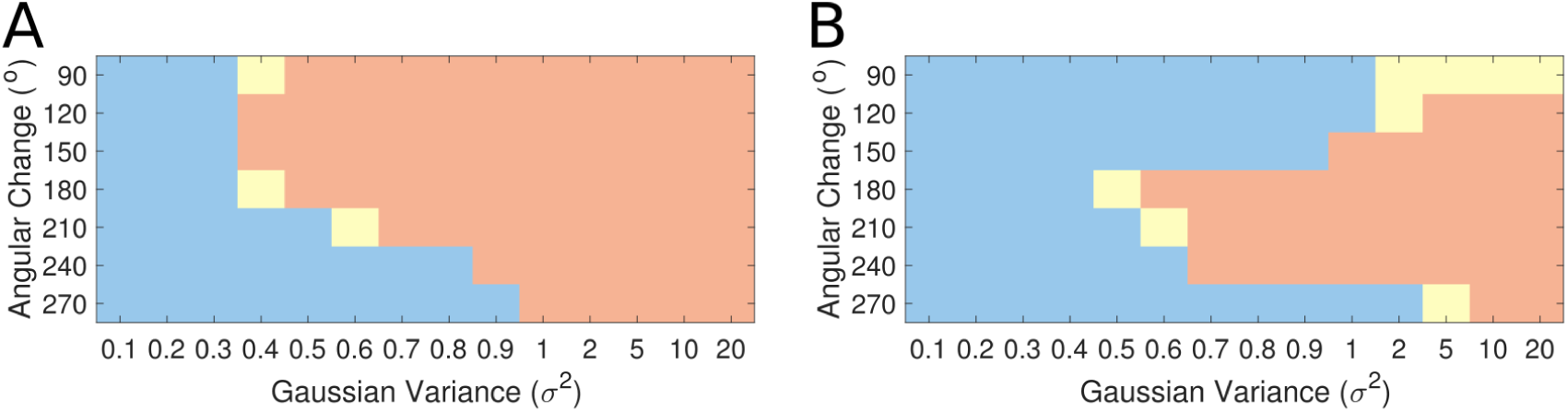
Transition regime as function of inhibitory uniformity. Heading signal transition regime for **A** the fruit fly ring attractor circuit and **B** the desert locust ring attractor circuit. Blue denotes gradual transition of the heading signal, orange denotes abrupt transition (jump), and yellow marks a mixture of the two regimes.

However, the morphology of the Delta7 neurons is not the only difference between the ring attractors in the two species, hence the recorded response patterns are not identical for the two species (***Figure 9***). There is anatomical difference in the presence of the P-EN to E-PG feedback loops only in the locust and consequently the synaptic weights differ between the two models. We investigate the effect of this anatomical difference in subsequent section.

### Attractor states distribution

We next investigated the fixed points of each ring attractor, that is, the attractor states. We stimulated the E-PG neurons by applying a von Mises spatially distributed stimulus with varying azimuthal centre around the attractor circuit. Both the fruit fly and locust circuits had eight discrete attractor states where the heading signal eventually settled once the stimulus was removed. Typically, the heading signal moved to the nearest attractor state. When a stimulus was applied equidistantly from two attractor states then, once the stimulus was removed, the ‘bump’ moved to one of the two stochastically (***Figure 10***). Ideally, if there was no noise, the heading signal would always move to the nearest attractor state which was not the case. The attractor states were more stable and clearly delineated in the locust while in the fruit fly there was a wider distribution of ‘bump’ locations, indicating that the locust ring attractor is more robust to drift and noise (***Figure 10***).

**Figure 10.**
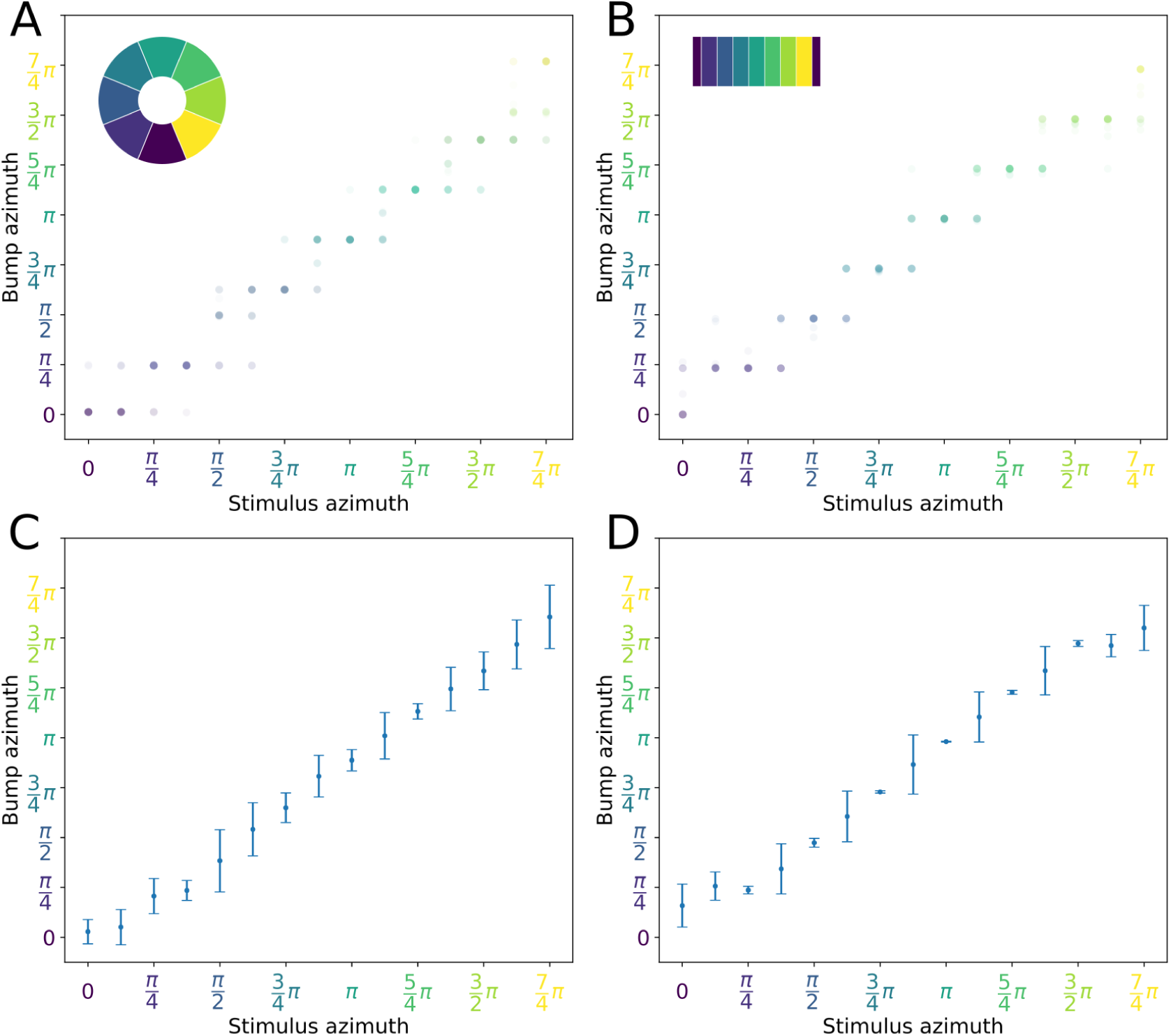
Distribution of activity ‘bump’ locations. Both ring attractor circuits have eight attractor states. However, the locust attractor states are more stable resulting to smaller dispersion of ‘bump’ location. On the abscissa (horizontal axis) the azimuth where von Mises stimulus is applied is shown. On the ordinate (vertical axis) the azimuth of the resulting activity ‘bump’ (attractor state), 3s after the stimulus is removed, is shown. Top row: The colour intensity of the discs indicates the frequency of each ‘bump’ location out of 40 trials for each stimulus. Discs are colour-coded by the ‘bump’ location 3s after stimulus removal. Inset images depict the corresponding EB tiles in colour. Bottom row: The mean location and standard deviation of the resulting activity ‘bump’. Smaller standard deviation corresponds to the ‘bump’ settling more frequently to the same azimuth. This is the case when the stimulus is applied near an attractor state. Applying stimulus equidistantly from two attractor states results to a movement of the ‘bump’ to either of them and hence the increased standard deviation. **A, C** is for the fruit fly **B, D** is for the locust. In locust when stimulating the ring attractor at one of the eight attractor states the ‘bump’ settles at it, indicated by the reduced standard deviation at these locations. In the fruit fly the activity ‘bump’ is prone to noise and not as stable, thus the standard deviation is not as modulated.

### Stability characteristics of the ring attractors

An important aspect of a ring attractor is its stability characteristics. The differences in the distribution of ‘bump’ locations reported in the previous section hinted that the locust ring attractor is more robust to noise. To quantify this property of the two ring attractors we measured the effect of different levels of structural (synaptic) noise to the circuit stability. The ring attractor of the locust was significantly more tolerant to structural noise than the fruit fly circuit (***Figure 11***A).

**Figure 11.**
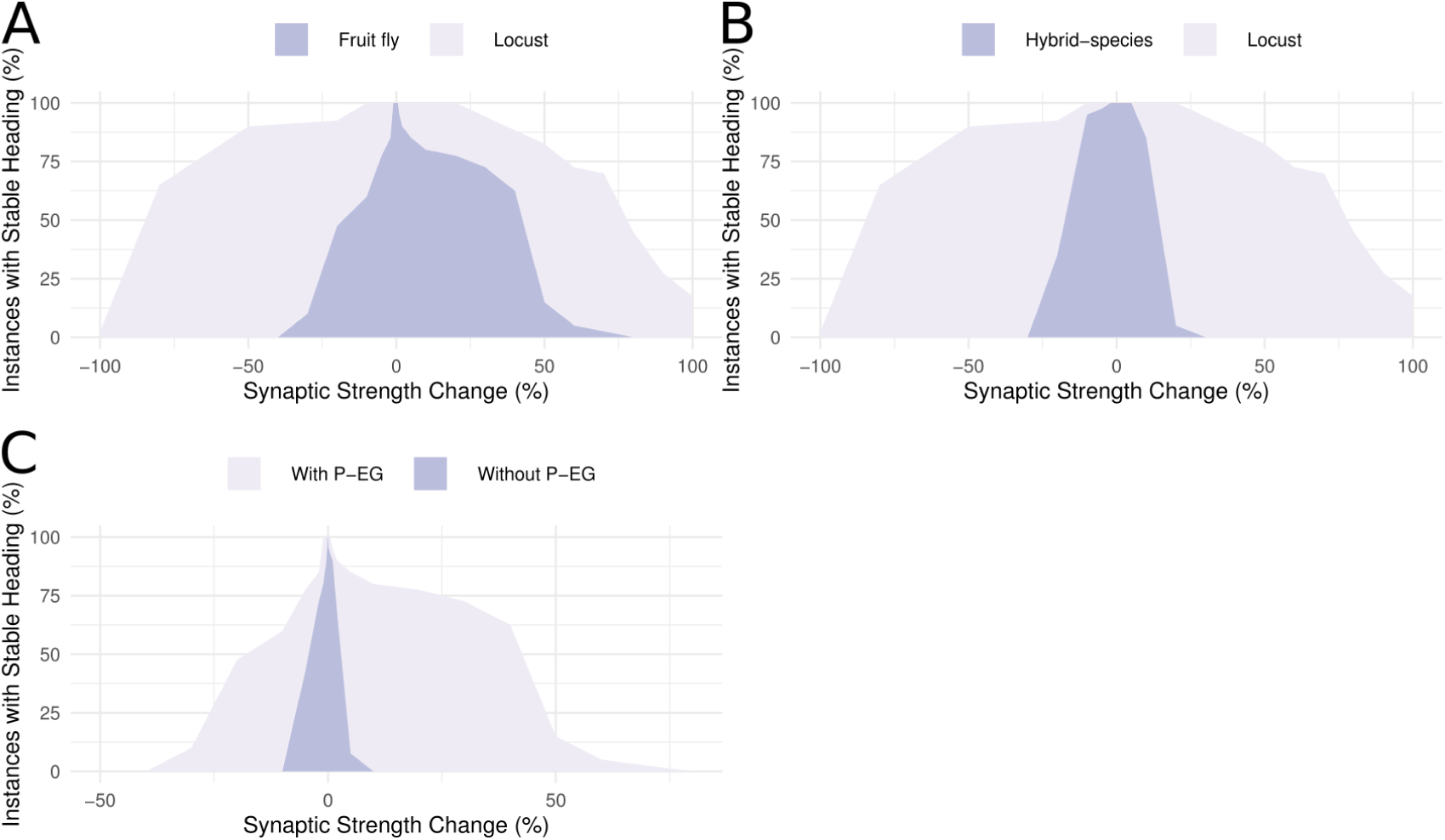
Ring attractor stability. **A** Stability of the ring attractor heading signal for the fruit fly and the locust model. **B** Stability of the ring attractor heading signal for the locust and hybrid-species (fruit fly with localised inhibition) model. Percentage of trials that result to a stable activity ‘bump’ as a function of structural asymmetry introduced by deviating synaptic strengths between P-EN and E-PG neurons (number of trials n=40 for each level of structural asymmetry). The locust ring attractor is more robust to such structural noise. **C** Stability of the fruit fly ring attractor heading signal when the circuit includes the P-EG neurons versus when they are removed (number of trials n=40 for each level of structural asymmetry). With P-EG neurons included the circuit maintains a stable heading signal for larger deviations of the synaptic strengths between P-EN to E-PG neurons.

However, these two ring attractors differ in several respects. To identify the reason for the reduced sensitivity of the locust model to synaptic noise we compared the locust with the hybrid-species model. These two models differ only in the existence of reciprocal connections between P-EN and E-PG neurons only in the locust. If these reciprocal connections are responsible for the increased robustness of the circuit we would expect the locust model to be more robust to synaptic noise than the hybrid-species model. This is exactly what we found (***Figure 11***B), thus we inferred that these reciprocal connections provide the increased robustness to the locust model. This circuit specialisation might have important repercussions to the behavioural repertoire of the species, enabling locusts to maintain their heading for longer stretches of time than fruit flies, an important competence for a migratory species such as the locust.

It is interesting to note that even though in the locust the E-PG to P-EG recurrency is weaker than in the fruit fly, the presence of the extra P-EN to E-PG recurrency in the locust results in a more stable ring attractor than that in the fruit fly which possesses only one but strong recurrency loop. Finally, the hybrid-species model is less robust than the fruit fly one. These models differ in the inhibitory domains distribution and their synaptic strengths. Even though this difference in robustness is smaller than the previously examined ones we can see an effect of the inhibitory pattern on the stability of the circuit.

### P-EG neurons stabilize the head direction circuit

In our models we included the P-EG neurons connecting the PB glomeruli with EB tiles. Unlike the P-ENs, these neurons have the same connectivity pattern as the E-PG neurons but with presynaptic and postsynaptic terminals on opposite ends. What is the effect of the P-EG neurons in the circuit? Effectively, the P-EG neurons form secondary positive feedback loops within each pair of columns that, we hypothesised, help the heading signal to be maintained stably in the current column, even when lacking external input. Therefore, we expected the circuit to function as a ring attractor without these connections, but to be more vulnerable to drift if the neuronal connection weights are not perfectly balanced. The recurrent P-EG to E-PG loop should counteract this tendency to drift.

To investigate the role of these neurons in the circuit, we measured the effect of imposing imbalance in the connectivity strengths of P-EN to E-PG neurons between the two hemispheres. We did this for the full fruit fly circuit and an altered circuit with the P-EG neurons removed. The synaptic strengths for the two circuits were optimised separately, since completely removing the P-EG neurons without appropriate synaptic strength adjustment breaks the ring attractor. We measured the percentage of simulation runs that resulted in a stable heading signal being maintained for at least 3 seconds. The presence of the P-EG neurons substantially increased the robustness of the circuit to the effects of imbalance of the P-EN to E-PG synaptic strength, as a stable heading signal was observed over a far wider range of weight changes (***Figure 11***C). The P-EG neurons therefore contribute significantly to the tolerance of the ring attractor to synaptic strength asymmetries.

### Response to proprioceptive stimuli

Mechanistically, Turner-Evans *et al.* showed that the activity of P-EN neurons in one hemisphere of the brain increases when the animal turns contralaterally, both with and without visual input (***Turner-Evans et al., 2017***). The increase in activity is related to the angular velocity the fly experiences (***Turner-Evans et al., 2017***). Whereas the source of the angular velocity information in darkness is not known, proprioception is the most likely sensory channel that can provide information about the rotational velocity the fly experiences. To test whether our models reproduce this behaviour, we artificially stimulated the P-EN neurons in one hemisphere of the PB, mimicking an angular velocity signal caused by turning of the animal, and observed the effect on the heading signal (***Figure 12***).

**Figure 12.**
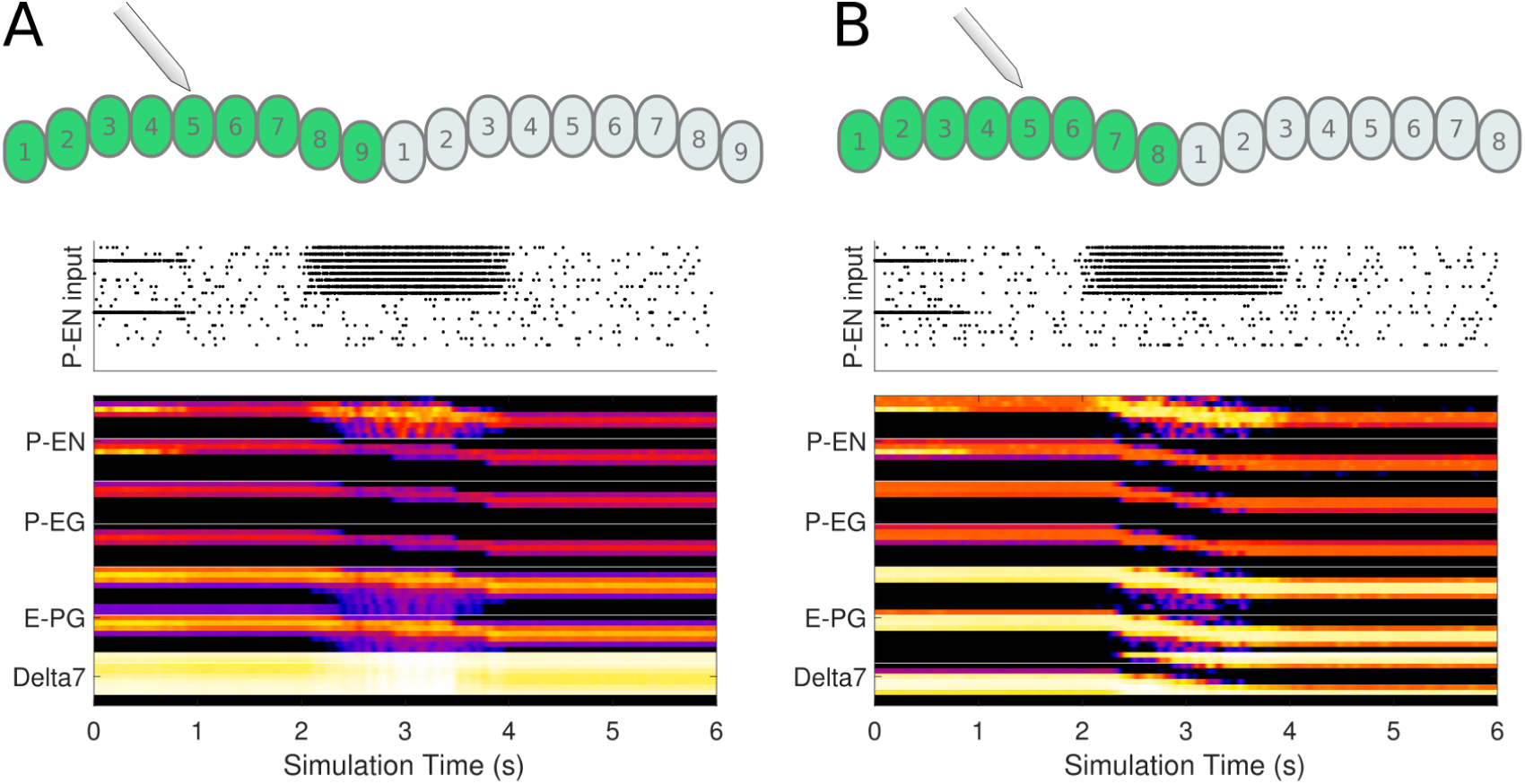
Response to uni-hemispheric stimulation. Upper plots show the P-EN stimulation protocol and corresponding induced P-EN activity; lower plots show the response of the ring attractor for **A** the fruit fly circuit and **B** the locust circuit. The initial bilateral stimulation initialises a persistent activity ‘bump’, which moves around the circuit in response to stimulation of P-EN in all the columns in one hemisphere only.

Both the locust and the fruit fly model reproduced the behaviour reported by ***Turner-Evans et al***. (***2017***). Exploration of the response of the circuit to different stimulation strengths showed that the rate by which the heading signal shifts around the ring attractor increases exponentially with increase of uni-hemispheric stimulation strength (***Figure 13***). While this general relationship was consistent between the two species, the increase was much steeper in the fly. Additionally, the required stimulus for initiating shifting is lower in the fruit fly. Both of these aspects concur with the faster response rate of the fruit fly ring attractor to positional stimuli and support the possibility to track fast saccades even when only angular velocity input is available.

**Figure 13.**
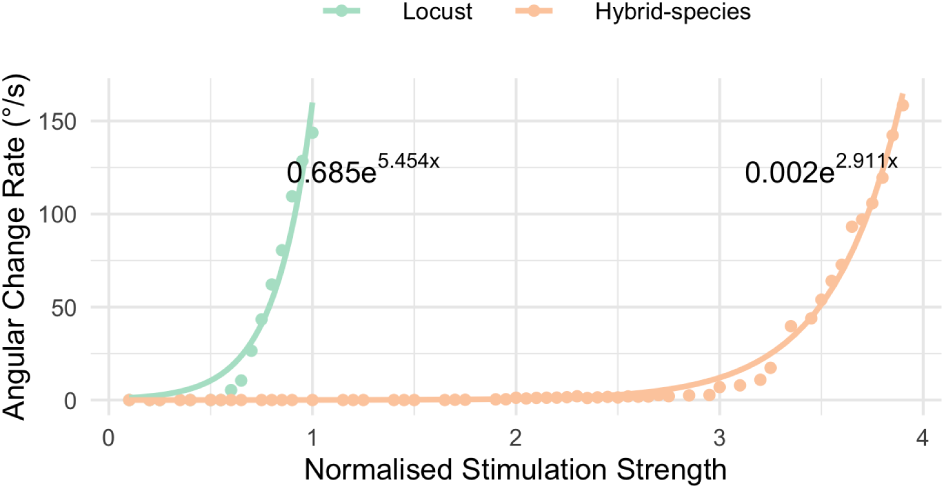
Response to uni-hemispheric stimulation. Response rate of change of the heading signal with uni-hemispheric stimulation of P-EN neurons. The angular rate of change increases exponentially with stimulation strength and does so most rapidly for the fruit fly circuit. The data points have been fit with the function *y* = *ae*^*bx*^ and the parameters of the fitted curves are shown on the plot.

Continuous application of rotational input causes the heading signal to reach an edge of the PB and then wraps around and continues on the other edge. This behavior is present in both models and is thus independent of the physical shape of the EB, i.e. whether it forms a closed ring or possesses open ends. The wrapping around of the heading signal is required for the animals to track movements that involve turning around its body axis for more that 360° and is supported by the effective eight-fold radial symmetry we found in both species.

## Discussion

The availability of tools for the study of insect brains at the single neuron level has opened the way to deciphering the neuronal organisation and principles of the neural circuit’s underlying behaviour. However, even where there is progress towards a complete connectome, the lack of data on synaptic strengths, transmitter identity, neuronal conductances, etc. leave many parameters of the circuit unspecified. Exploring these parameters via computational modeling can help to illuminate the functional significance of identified neural elements. We have applied this approach to gain greater insight into the nature of the heading encoding circuit in the insect central complex (CX), including the consequences of differences in circuit connectivity across two different insect species.

### Overall conservation of structure and function

We have focused on a subset of neurons in the PB and EB which have been hypothesised to operate as a ring attractor, with a ‘bump’ of neural activity moving across columns consistently with the changing heading direction of the animal. The neuronal projection patterns and columnar organisation differ between the two insects we have analysed, the fruit fly and the locust. There are additional morphological columns in the PB of flies (9 vs. 8), resulting in a different number of functional units that could in2uence the symmetry of underlying neural circuits. Also, the EB in the fruit fly forms a physical ring, while the homologous region in the locust is an open structure. Our analysis of the connectivity as a directed graph has revealed, surprisingly, that the circuits are nevertheless equivalent in their effective structure, forming a closed loop ring attractor with eight-fold symmetry in both species, with an identical functional role for each neuron class. The stability of this circuit across 400 million years of evolutionary divergence suggests that it is an essential, potentially fundamental, part of the insect brain.

### Differences in dynamical response

Two remaining, apparently more subtle, differences in the morphology between the two species were shown to have significant effects on the dynamical response of the heading direction circuit. First, the shape of the dendritic arborizations of one type of CX neuron determined how fast the model circuit could track rotational movements. Hence, a seemingly subtle change in morphology caused not only a change in response dynamics, but in fact, it altered the type of ring-attractor implemented by the circuit. Second, a difference in the overlap of neuronal projections in the EB results in an extra feedback loop between the P-EN and E-PG neurons in the locust circuit that makes it more robust to synaptic noise.

We suggest that the effect of these differences are consistent with the behavioural ecology of the two species. On the one hand, the faster response of the ring attractor circuit in the fruit fly accommodates the fast body saccades that fruit flies are known to perform (***Tammero and Dickinson, 2002***; ***Fry et al., 2003***). On the other hand, the locust is a migratory species, a behaviour that demands maintenance of a defined heading for a long period of time. This requirement for stability might have provided the selective pressure needed to drive the evolution of a more noise resilient head direction circuit (***de Vries et al., 2017***).

Although the ring attractors in the two species are substantially similar, the identified specialisations have a significant effect on their dynamics, thus raising the question of how valid generalisations drawn from studies in any one species can be. This is particularly relevant as most recent data have resulted almost exclusively from work in *Drosophila*.

### Assumptions and simplifications

As any model, our circuits are necessarily condensed and simplified versions of the real circuits in the insect brain. In comparison to previous models, the work we present has been more precisely constrained by the latest anatomical evidence and uses realistic biophysical properties, with realistic background activity to produce realistic spiking rates. Furthermore, in building our models we did not assume that the underlying circuit must be a ring attractor, but rather asked and investigated whether, given the available connectivity data, it can be. This was especially the case for the locust model since our work represents the first model of this circuit to date. Nevertheless it is important to outline those areas where our assumptions cannot be fully justified from the existing data, and the potential consequences for the model results.

#### Morphological Assumptions

In our model of the fruit fly heading tracking circuit, we assumed a uniform distribution of dendrites across the Delta7 neurons. Imaging of these neurons suggests that there might be a subtle variation of the dendritic density along their length. However, it is unclear how this subtle variation might be related to the synaptic density and efficacy. We, therefore, made the simplifying assumption and modeled these neurons as having uniform synaptic efficacy across the PB. However, we also explored the effect of varying the degree of uniformity, showing that there is a range of distributions that still can produce the fly-like rapidity in the circuit response.

In general, arborisation trees of neurons in the CX can be very complex, as they are not only confined to specific slices, but also to one or several layers, especially within the EB. In *Drosophila*, the spiny terminal arbors of E-PG neurons extend to the width of single wedges in the EB, occupying both the posterior and medial layers. In contrast, P-EG and P-EN neurons arborise in tiles, hence innervating only the posterior surface volume of the EB (***Wolff and Rubin, 2018***). Therefore, we assume that presynaptic terminals of P-EG and P-EN neurons form synapses with E-PG postsynaptic terminals in the posterior layer of the EB. In locusts, the E-PG arborizations are more complex, as these cells innervate a single wedge for anterior and medial EB layers, but extend at least twice this width to either side in the posterior layer that provides overlap with the P-EN neurons (***Heinze and Homberg, 2008***). Additionally, the wider fibers have a different morphological appearance. P-EG neurons in this species innervate all layers evenly. Although these detailed differences likely have consequences for connectivity, we simplified these arborisations to their most essential components, aiding the extraction of core features. With the advance of comparative connectomics, these aspects will become accessible for investigation.

#### Connectivity Assumptions

Several assumptions were made while deriving the neuronal connectivity in our models. We assumed well delineated borders of synaptic domains which is clearly not always the case, especially in the EB as some overlapping of synaptic domains due to stray terminals is to be expected (***Wolff et al., 2015***). The circumferential extent of arbors in wedges and tiles may affect the integrity of the resulting circuit and its properties. However, due to lack of adequate data about the extent of such overlap we cannot currently model this aspect in a sensible way.

Furthermore, neuronal connectivity was mostly inferred by co-location of neuron arbors, that is projection patterns. A functional connectivity study has reported that stimulation of E-PG neurons triggered significant responses to Delta7 neurons but no columnar neurons (***Franconville et al., 2018***). However, as those authors note, the lack of response might be due to the limitations of the method used. Alternatively, such connections might be mediated by interneurons instead of being monosynaptic. Future work using electron microscopy data will elucidate which of the overlapping arborisations correspond to functional connections and allow us to augment our models.

#### Functional Assumptions

Further assumptions were made about the neuron polarity, type of synapses and synaptic efficiencies. ***Lin et al.*** (***2013***) characterise the EB arbor of E-PG neurons in *Drosophila* as having both presynaptic and postsynaptic domains, however, ***Wolff et al.*** (***2015***) report that using antisynaptotagamin is inconclusive for presynaptic terminals. In our models for both the fruit fly and the locust we thus assume E-PG neurons are purely postsynaptic in the EB, following the most conservative polarity estimate.

Furthermore, the Delta7 neurons are assumed to have inhibitory effect to their postsynaptic neurons, as ***Kakaria and de Bivort*** (***2017***) proposed. However, there is some evidence that Delta7 neurons possibly make both inhibitory and excitatory synapses to other neurons (***Franconville et al., 2018***). As those neurons (P-FN neurons) are not part of our current model, we make the simplifying assumption that Delta7 neurons have exclusively inhibitory effect on their postsynaptic neurons. Finally, it is possible that there are other sources of inhibition in the circuit, e.g. mediated by the ring neurons as suggested by ***Green and Maimon*** (***2018***). We do not explore this possibility in our current work.

We assumed that the synaptic strength of all synapses of each class are identical. This might not be the case in the actual animals, especially considering that one of the EB tiles (T1) is innervated by twice as many neurons as other tiles (Supplementary Fig. S2). Neurons innervating this tile might have reduced synaptic efficacy in order to maintain the radial symmetry of the circuit intact. Similarly, synaptic strength variation might exist for the two Delta7 neurons that have presynaptic terminals in three glomeruli instead of two.

#### Biophysical Assumptions

All types of neurons in our models are assumed to have the same biophysical properties even though anatomical evidence has shown that their morphology, somata size and main neurite thickness differ (***Heinze and Homberg, 2008***). We used a point spiking neuron model, which was sufficient for investigating the performance characteristics of the ring attractors when exposed to realistic neuronal noise, but clearly is highly abstracted relative to real neurons. However, the use of multicompartmental neural models would have increased the complexity without offering tangible gains, as most of the necessary detail to constrain such models is still unknown.

### Comparison to ‘canonical’ ring attractor models

In our work we compared the hypothetical heading tracking circuit of two evolutionary distant species. We went beyond mere simulation of neuronal projection data by analysing and deriving the effective underlying circuit structure of the two ring attractors. Our analysis and derivation of the complete effective neuronal circuits revealed not only differences in dynamics but also the construction principles of these evolved ring attractor circuits. This approach allowed us to identify elements that differ in several ways from the ‘canonical’ ring attractor as described in earlier theoretical models (e.g. ***Amari*** (***1977***); ***Skaggs et al.*** (***1995***); ***Zhang*** (***1996***)).

For example, the circuit found in the two insects combines two functionalities in the P-EN neurons that are typically assigned to separate neural populations in computational models of ring attractors: one set of neurons to provide the lateral excitation of nearest neighbours; and a different set neurons that receive angular velocity input to drive the left-right rotation of the heading signal. In the insect circuit, the P-EN cells are part of the lateral excitation circuit, providing excitation to their two nearest neighbours, but also receive angular velocity input in the noduli. This difference is suggestive of a more efficient use of neuronal resources than the typical computational models of ring attractors. Another novel element we found in the insect ring attractors is the presence of local feedback loops within each octant of the circuit structure (P-EG to E-PG and P-EN to E-PG). We found that both of these feedback loops increase the tolerance of the ring attractors to noise.

### Hypotheses regarding circuit differences

Another unique aspect of our model is the comparison of related, but not identical, circuits between two species. Indeed, using computational modelling allows us to investigate ‘hybrid’ circuits, combining features of each, to try to understand the functional significance of each observed difference independently. Nevertheless, some differences between these circuits are not explained by the current model, and may require additional work to fully explicate.

One question is what is the role, if any, of the ninth PB glomeruli found so far only in *Drosophila*? In particular, the existence of the innermost glomeruli that are not innervated by the P-EN neurons seems perplexing. The same signals from tile 1 of the EB are sent to both ends of each hemisphere of the PB (glomeruli 1 and 9) and from there action potentials propagate along the Delta7 neurons along the PB length. Our speculation is that this may be a mechanism to reduce the distance and time these signals have to travel to cover the full PB, i.e., the maximum distance any signal must travel is only half of distance as it would be to propagate from one end of the PB to the other as in other species, such as the locust. If this is the case, it would constitute one more specialisation in *Drosophila* that reduces the response time of the ring attractor. It therefore seems that several specialisations have been orchestrated in minimising the response delays in fruit flies. Testing this idea would require multicompartmental models to capture the action potential transmission time along neurites; as argued above, this may need preceding detailed biophysical characterisation of the Delta7 neurons.

Another remaining question is what is the role of the closed ring shaped EB in *Drosophila*. One possibility is that such a closed ring topology would allow reciprocal connections between P-EN and E-PG neurons. This would allow direct propagation of signals between these neurons within the EB instead of requiring them to travel via the PB as in the current model, and might again increase the speed with which the heading direction can be tracked, and allow smoother transition between neighbouring tiles. Note that such reciprocal connections within the EB can only be continuous with a closed ring anatomy and would not be possible between the two ends of the EB in the locust. To investigate the existence of such hypothetical reciprocal connections within the EB, further electron microscopy neurobiological studies are required. Possibly blocking signal transmission via the PB to isolate functional connectivity within the EB would reveal if there are such reciprocal connections. Subsequently, signal transmission time measurements within the EB versus via the PB would determine how different and hence significant those two pathways might be in the ring attractor performance.

A further hypothesis relates to the evolutionary lineage of these two features in the *Drosophila* CX. It will be of interest to study whether the ring shaped EB appeared before or after the appearance of the ninth glomeruli. One possibility is that the EB evolved into a ring shape after the appearance of the ninth glomeruli in the PB, allowing connections from one common tile to both glomeruli 1 and 9 and hence providing such a common driving signal. Alternatively, a pre-existing ring shaped EB might have allowed the evolution of usable ninth glomeruli that resulted in faster propagation. Similarly, the P-EN to E-PG recurrency found only in the locust might be an acquired adaptation of the locust that increases robustness to noise, or a common feature among other insect species that has been lost by fruit flies.

Comparison of different species could potentially elucidate such questions. We would expect individual species to have a selective subset of the specialisations we found, endowing them with brain circuits supporting the behavioural repertoire suiting their ecological niche. It will, therefore, be informative to analyse the effective heading direction circuit of other species, spanning evolutionary history, in order to get insights into how such adaptations relate to and accommodate behaviour. Our results emphasise the importance of comparative studies if we are to derive general principles about neuronal processing, even in systems that appear highly conserved such as the CX head direction circuit in insects. Many of the circuit properties observed in *Drosophila* appear to reflect specific evolutionary adaptations related to tracking rapid flight maneuvres. Despite the many strengths of *Drosophila* as an experimental model, it remains important to ground conclusions about the insect brain in comparison with other species.

## Methods and Materials

### Neuron model

Our models are based on the source code published by ***Kakaria and de Bivort (2017)***. We used Leaky Integrate and Fire neuron models (***Stein, 1967***). The membrane potential of each neuron was modelled by the differential equation

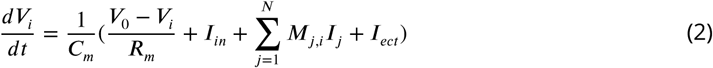

where *V*_*i*_ is the membrane Voltage of neuron *i, I*_*in*_ the input current, *M*_*j,i*_ the network connectivity matrix, *I*_*j*_ the output current of each neuron in the circuit, and *I*_*ect*_ is the ectopic current produced by optogenetic or thermogenetic manipulation.

The model parameter values including membrane resistance, capacitance, resting potential, undershoot potential and postsynaptic current magnitude (*I*_*PSC*_) and delay were set to the same values as used by ***Kakaria and de Bivort*** (***2017***). These values are consistent with evidence from measurements in *Drosophila melanogaster* and other species. The membrane capacitance, *C*_*m*_, is set to 0.002*µF* for all neurons assuming a surface area of 10^−3^*cm*^2^ and the membrane resistance *R*_*m*_ to 10*M*Q (***Gouwens and Wilson, 2009***). The resting potential *V*_0_ is set to −52*mV* for all neurons (***Rohrbough and Broadie, 2002***; ***Sheeba et al., 2008***). The action potential threshold is −45*mV* (***Sheeba et al., 2008***; ***Gouwens and Wilson, 2009***). When the membrane voltage reaches the threshold voltage an action potential template is inserted in the recorded time series. The action potential template is defined as (***Kakaria and de Bivort, 2017***):

The action potential template is defined as

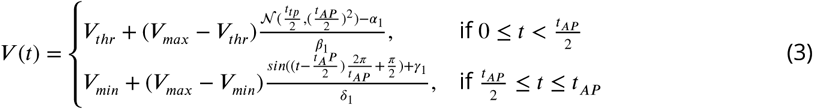

where *V*_*max*_ is the peak voltage set to 20*mV* (***Rohrbough and Broadie, 2002***). *V*_*min*_ is the action potential undershoot voltage, set to −72*mV* (***Nagel et al., 2015***). *t*_*AP*_ is the duration of the action potential set to 2ms (***Gouwens and Wilson, 2009***; ***Gaudry et al., 2012***). 𝒩 (*µ, σ*^2^) is the probability density function of a Gaussian with a mean *µ* and standard deviation *σ*^2^. *α*_1_, *β*_1_, *γ*_1_, and *δ*_1_ are normalisation parameters for scaling the range of the Gaussian and the sinusoidal to 0 to 1.

The firing of an action potential also adds a postsynaptic current template to the current time series. The postsynaptic current template is defined as

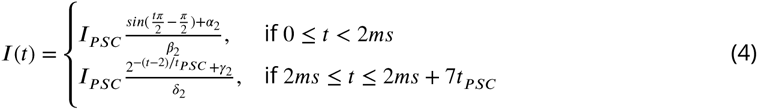

where *I*_*PSC*_ = 5*nA* (***Gaudry et al., 2012***). Excitatory and inhibitory postsynaptic currents are assumed to have the same magnitude but opposite signs. *t*_*PSC*_ = 5*ms* is the half-life of the postsynaptic current decay (***Gaudry et al., 2012***). *α*_2_, *β*_2_, *γ*_2_, and *δ*_2_ are normalisation constants so that the range of the sinusoidal and exponential terms is 0 to 1. The postsynaptic current traces have duration 2*ms* + 7*t_PSC_*, that is 2*ms* of rise time plus 7*t_PSC_* of decay time. The simulation was implemented using Euler’s method with a simulation time step of 10^−4^*s*. Our code is derived from the source code published by ***Kakaria and de Bivort (2017)***.

### Neuronal projections and connectivity

We modeled and compared the hypothetical ring attractor circuits of the fruit fly *Drosophila melanogaster* and the desert locust *Schistocerca gregaria*. The connectivity of the circuits has been inferred mostly from anatomical data derived using light microscopy, with overlapping neuronal terminals assumed to have synapses between them (***Wolff and Rubin, 2018***; ***Wolff et al., 2015***; ***Heinze and Homberg, 2007, 2008***; ***Pfeiffer and Homberg, 2013***).

Our models include the E-PG, P-EG, P-EN and Delta7 neurons. Note that P-EG refers to the updated set of neurons innervating all PB glomeruli as reported in ***Wolff and Rubin*** (***2018***) (PBG1–9.s-EBt.b-D/V GA.b). In this paper E-PG refers to the E-PG (PBG1–8.b-EBw.s-D/V GA.b) and the complimentary E-PG_T_ (PBG9.b-EB.P.s-GA-t.b) combined (***Wolff et al., 2015***; ***Wolff and Rubin, 2018***). Therefore, E-PG neurons are innervating all PB glomeruli. Delta7 refers to PB18.s-GxΔ7Gy.b and PB18.s-9i1i8c.b neurons combined (***Wolff et al., 2015***; ***Wolff and Rubin, 2018***). ***Table 3*** shows this correspondence in detail.

These neurons innervate two of the Central Complex neuropils, the Protocerebral Bridge (PB) and the Ellipsoid Body (EB). Ellipsoid Body is the name used for this structure in the fruit fly *Drosophila melanogaster* while in the locust *Schistocerca gregaria* the equivalent structure is referred to as lower division of the central body (CBL). To aid comparisons with previous models and for general simplification, we will use the term EB for both species. The PB is a moustache shaped structure consisting of 16 or 18 glomeruli, depending on the species. In the fruit fly *Drosophila melanogaster* the EB has a torus shape consisting of 8 tiles. Each tile is further broken down in two wedges. In the locust *Schistocerca gregaria* the EB (CBL) is a linear structure, open at the edges, consisting of 8 columns. Each column has two subsections similar to the wedges found in *Drosophila melanogaster*.

For both *Drosophila melanogaster* and *Schistocerca gregaria*, the synaptic domains of each of the E-PG, P-EN and P-EG neurons are confined to one glomerulus of the PB, ***Figure 3***. In the EB the synaptic domains of E-PG neurons are constrained in single wedges (half tiles) while the synaptic domains of P-EN and P-EG neurons extend to whole tiles (***Wolff et al., 2015***).

Furthermore, E-PG neurons innervate wedges filling the posterior and medial shells of the EB while P-EG neurons innervate whole tiles filling only the posterior shell of the EB (***Wolff et al., 2015***). Our model assumes that their overlap in the posterior shell equals functional connectivity. Finally, P-EN neurons innervate whole tiles of the EB (***Wolff et al., 2015***).

In our models, the E-PG, P-EG and P-EN neurons are assumed to produce excitatory effect to their postsynaptic neurons while Delta7 neurons are assumed to provide the inhibition, as ***Kakaria and de Bivort*** (***2017***) proposed. The projection patterns of the aforementioned neurons were mapped to one connectivity matrix for each species (***Figure 1***). ***Figure 1***G shows the connectivity matrix of the *Drosophila melanogaster* fruit fly model, ***Figure 1***H the connectivity matrix of the the *Schistocerca gregaria* desert locust model.

The most salient difference between the two matrices is the connectivity pattern of the Delta7 neurons (lower right part of 1G and 1H). In *Drosophila melanogaster* the Delta7 neurons receive and make synapses uniformly across the PB glomeruli while in the locust *Schistocerca gregaria* the Delta7 neurons have synaptic domains focused in specific glomeruli. We analysed the effect of this difference in detail in the Results section. Another major difference apparent in the connectivity matrices is the existence of 18 glomeruli in the PB of *Drosophila melanogaster* but 16 in *Schistocerca gregaria*.

We modeled each PB glomerulus, as being innervated by one neuron of each class (E-PG, P-EG, P-EN) even though in reality there are several copies of each one.

The locust inhibition has been modeled as a summation of two Gaussian functions that approximate the synaptic density across the PB glomeruli, estimated by visual inspection of light microscopy data. The variance (*σ*^2^) of the Gaussian functions was set to the value 0.4 as the nearest approximation to the visually determined synaptic domain width.

In all our simulations we use the full connectivity matrices derived from neuronal projection patterns data and not the effective circuits described here.

### Stimuli

Two types of input stimuli are provided to the circuit: heading and angular velocity. The heading stimulus is provided as incoming spiking activity directly to the E-PG neurons, corresponding to input from Ring neurons (***Young and Armstrong, 2010***) (called TL-neuron in other species (***Vitzthum et al., 2002***)). The position of a visual cue, angle of light polarization (***Heinze and Homberg, 2007***) or retinotopic landmark position (***Seelig and Jayaraman, 2015***) around the animal, is mapped to higher firing rates of E-PG neurons in the corresponding wedge of the EB. We assumed that the background activity of the upstream Ring neurons is produced by a Poisson process with a mean action potential rate of 5 impulses/s. The peak impulse firing rate of the stimulus signal was equal to the peak spiking rate of the activity ‘bump’ of the E-PG neurons under steady state conditions in order to have comparable measurements across experiments. These spiking trains are also produced by a Poisson process. The angular velocity stimulus consists of spikes which are directly supplied to all P-EN neurons in one hemisphere of the PB, corresponding to the direction of motion. The peak impulse rate of the injected spike trains was equal to the peak rate of the steady state activity ‘bump’ across the P-EN neurons. This was done in order to allow for direct comparisons between experiments. Clockwise rotations of the animal were passed to the left/right side of the PB and vice versa for counter-clockwise rotations.

### Free parameters

The free parameters of our models are the synaptic strengths. The synaptic strengths of synapses connecting each class of neurons are assumed to be identical, e.g, all P-EN to E-PG synapses have the same strength. Therefore, we have one free parameter for each synaptic class. Furthermore, we reduced the complexity of the problem by making the synaptic strength between some classes of neurons identical. The synaptic strengths of E-PG to P-EN and P-EG are identical as are the synaptic strengths of Delta7 to P-EN and P-EG. We used the minimum set of synaptic strengths that result in a working ring attractor. We assumed that all synapses are excitatory apart from the synapses with Delta7 neurons on the presynaptic side, which are assumed to be inhibitory, as ***Kakaria and de Bivort*** (***2017***) proposed. The synaptic strength was modeled as the number of *I*_*PSC*_ unit equivalents flowing to the postsynaptic neuron per action potential.

Whereas our models are constrained by anatomical evidence, existing biological studies do not specify synaptic weights or connectivity in general with the exception of ***Franconville et al.*** (***2018***). Based on the goal that each of the circuits should yield a functional ring attractor, an optimization algorithm was used to search for synaptic strength combinations that result in working ring attractors. Both simulated annealing and particle swarm optimization algorithms were used; the first one converges quicker while the second one covers the search space more thoroughly. We constrained the acceptable solutions to those that produced an activity ‘bump’ with full width at half maximum (FWHM) of approximately 90° since this is the width that has been observed in fruit flies (***Kim et al., 2017***).

The optimisation algorithm was run to optimise the synaptic separately for each of the models: the *Drosophila*, the locust, and the hybrid-species model. All source code will be made available on github.

## Supplemental information

**Fig. S1.**
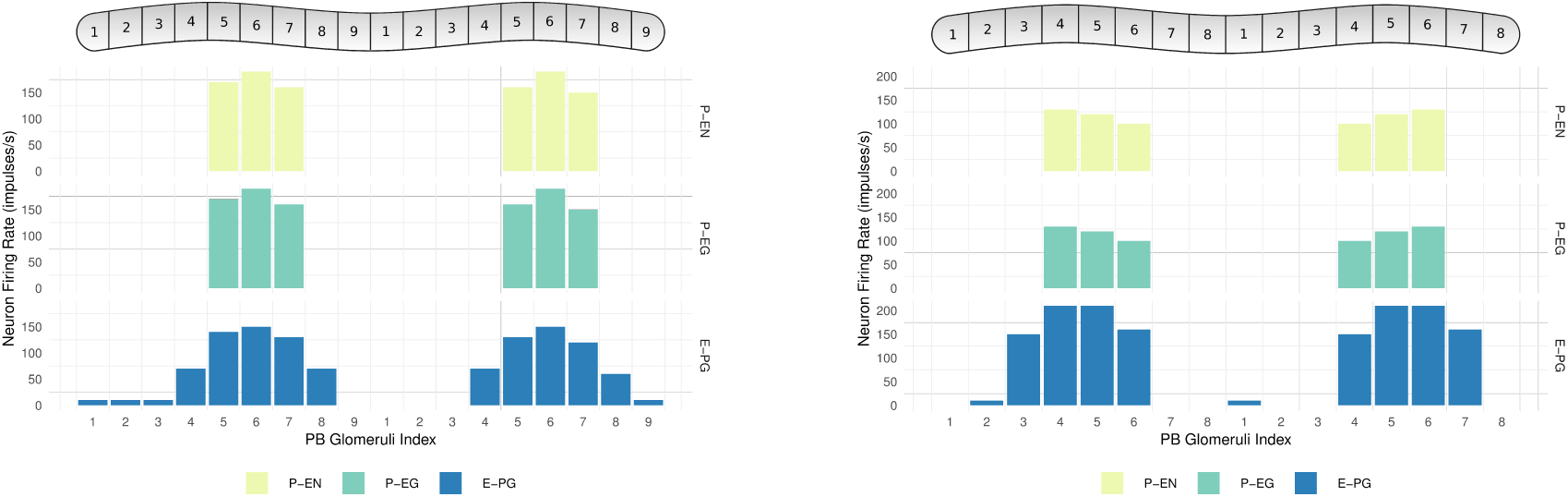
Neuronal activity across PB glomeruli. Figures (a) and (b) show the neuronal activity of P-EN, P-EG and E-PG neurons innervating the glomeruli of the PB for the simulated model of (a) *Drosophila melanogaster* and (b) *Schistocerca gregaria*. The activity ‘bump’ is centred around identically numbered glomeruli on the two hemispheres.

**Fig. S2.**
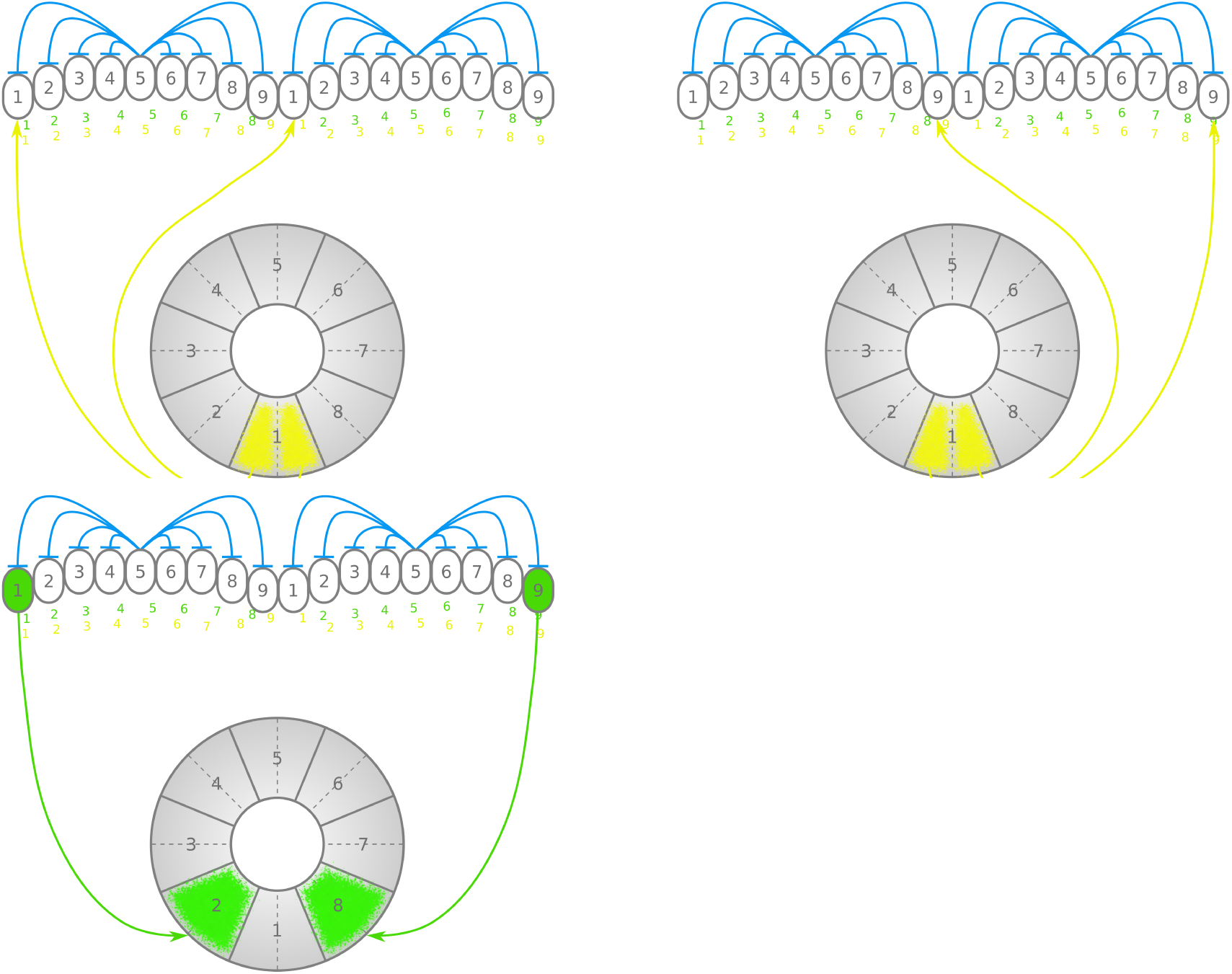
Neuronal connectivity of *Drosophila melanogaster*. Both the E-PG_1_ E-PG_9_ neurons innervate with postsynaptic terminals the same EB tile W1. However, only one PB glomerulus is innervated by P-EN neurons in each hemisphere, that is, G1 on one side and G9 on the other. This results in the same number of P-EN neurons innervating tiles W2 and W8 as any other tile, hence not breaking the symmetry of the ring attractor.

**Fig. S3.**
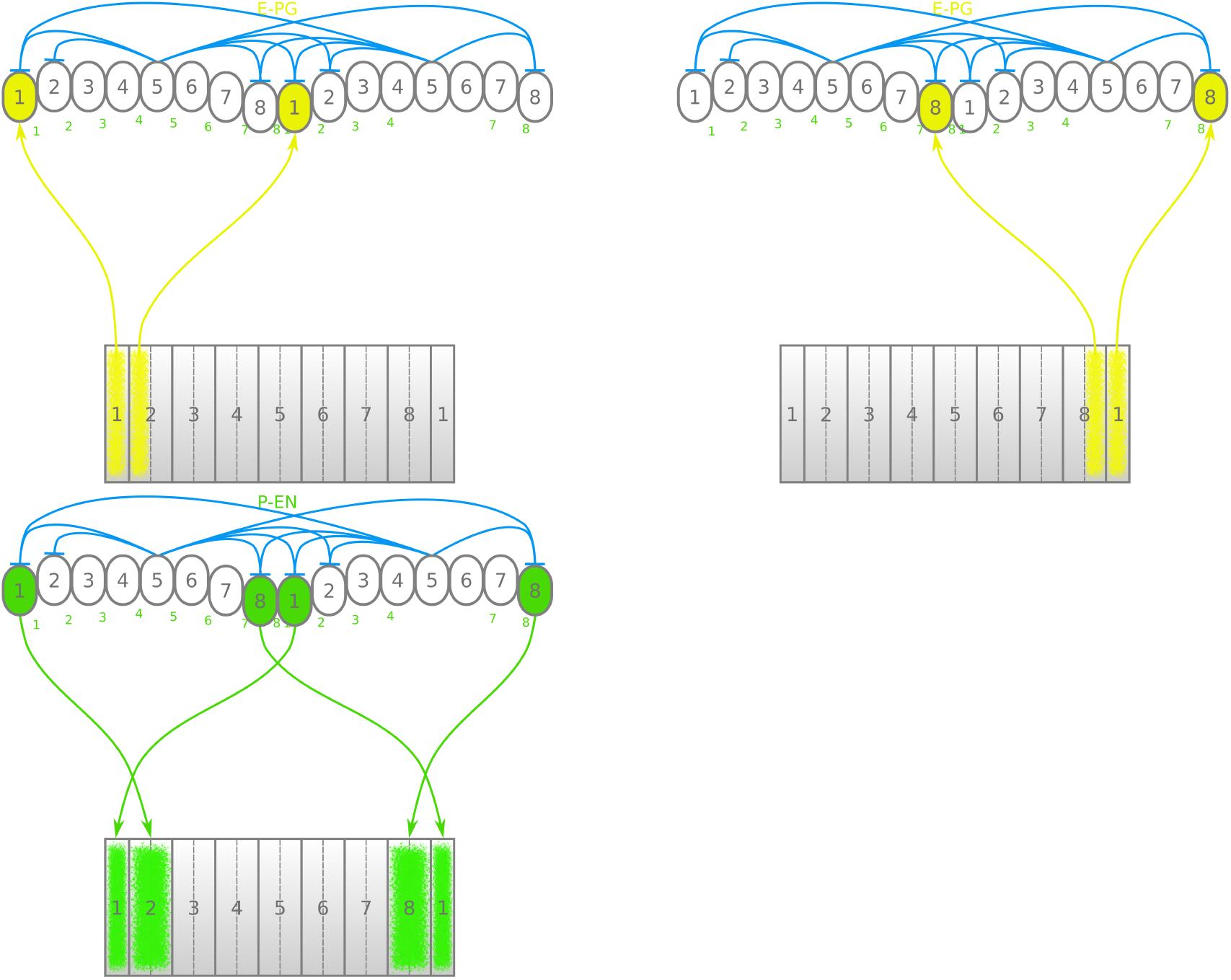
Neuronal connectivity of *Schistocerca gregaria*. The locust anatomy differs from the fruit fly in the number of PB glomeruli. Since in the locust there are as many glomeruli as EB columns the same number of E-PG and P-EN neurons innervate all glomeruli and columns maintaining the symmetry of the ring attractor.

## Acknowledgments

Financial support from the European Research Council (ERC) under the European Union’s Horizon 2020 research and innovation program (grant agreement no. 714599). The founders had no role in study design, data collection and interpretation, or the decision to submit the work for publication.

